# Adaptations of the pregnant murine heart to pre-existing hypertension

**DOI:** 10.64898/2026.09.14.751233

**Authors:** Elnaz Ghajar-Rahimi, Arden Shen, Adalyn Meeks, Molly Kaissar, Shaheer Faruqi, Samantha Stebbings, Sarah Grev, Jake Castro, Abigail Cox, Kyoko Yoshida, Craig J. Goergen

**Author notes:** Correspondence: Craig J. Goergen, 206 S. Martin Jischke Drive, West Lafayette, IN 47907.

## Abstract

2-5% of pregnancies are complicated by chronic hypertension [1], increasing the risk of peripartum cardiomyopathy and maternal morbidity [2, 3]. Hypertension reduces stroke volume, whereas pregnancy increases stroke volume. The combined impact of these opposing loads on cardiac function and strain remains unclear. We used a mouse model of pregnancy with pre-existing hypertension to characterize cardiac remodeling. We hypothesized that pregnancy may counteract hypertension-related reductions in cardiac function. Nulliparous female C57Bl6/J mice (7-11 weeks old) were assigned into pregnant (PREG, n=11), angiotensin II-treated (ANGII, n=7), and ANGII+PREG (n=8) groups. Hypertension was induced with subcutaneous angII infusion (1000ng/kg/min). Cardiac function was measured with high-frequency ultrasound. Ejection fraction (EF) was maintained in PREG animals at 65% but decreased significantly by day 36 in ANGII animals (44.85*±*3.27%). In ANGII+PREG mice, EF initially decreased but trended towards PREG values on postpartum day 1. Left ventricle strain was lower in ANGII and ANGII+PREG groups compared to PREG. ANGII and ANGII+PREG animals showed decreased early longitudinal diastolic strain rate at mid-pregnancy compared to baseline (0.82*±*0.09% ANGII; 0.73*±*0.09% ANGII+PREG), while increasing in PREG (1.76*±*0.14%). Minimal physiologically relevant fibrosis was noted across groups via immunohistochemistry. Heart mass-to-body mass was lowest in PREG (6.20*±*0.22mg/g), greatest in ANGII (8.95*±*0.39mg/g), and intermediate in ANGII+PREG (6.92*±*0.25mg/g). Cardiomyocyte cross-sectional area was similar between groups. Overall, pregnancy altered the functional and hypertrophic response to elevated blood pressure. Findings from this work set the foundation for future efforts that may eventually improve cardiovascular care for pregnant patients.

**New & Noteworthy:** 4D strain analysis reveals subtle cardiac decline in pregnant mice treated with angiotensin II, despite adaptations to global cardiac function and subdued hypertrophy. These findings suggest potential cardioprotective effects of pregnancy.

**Visual Abstract:** 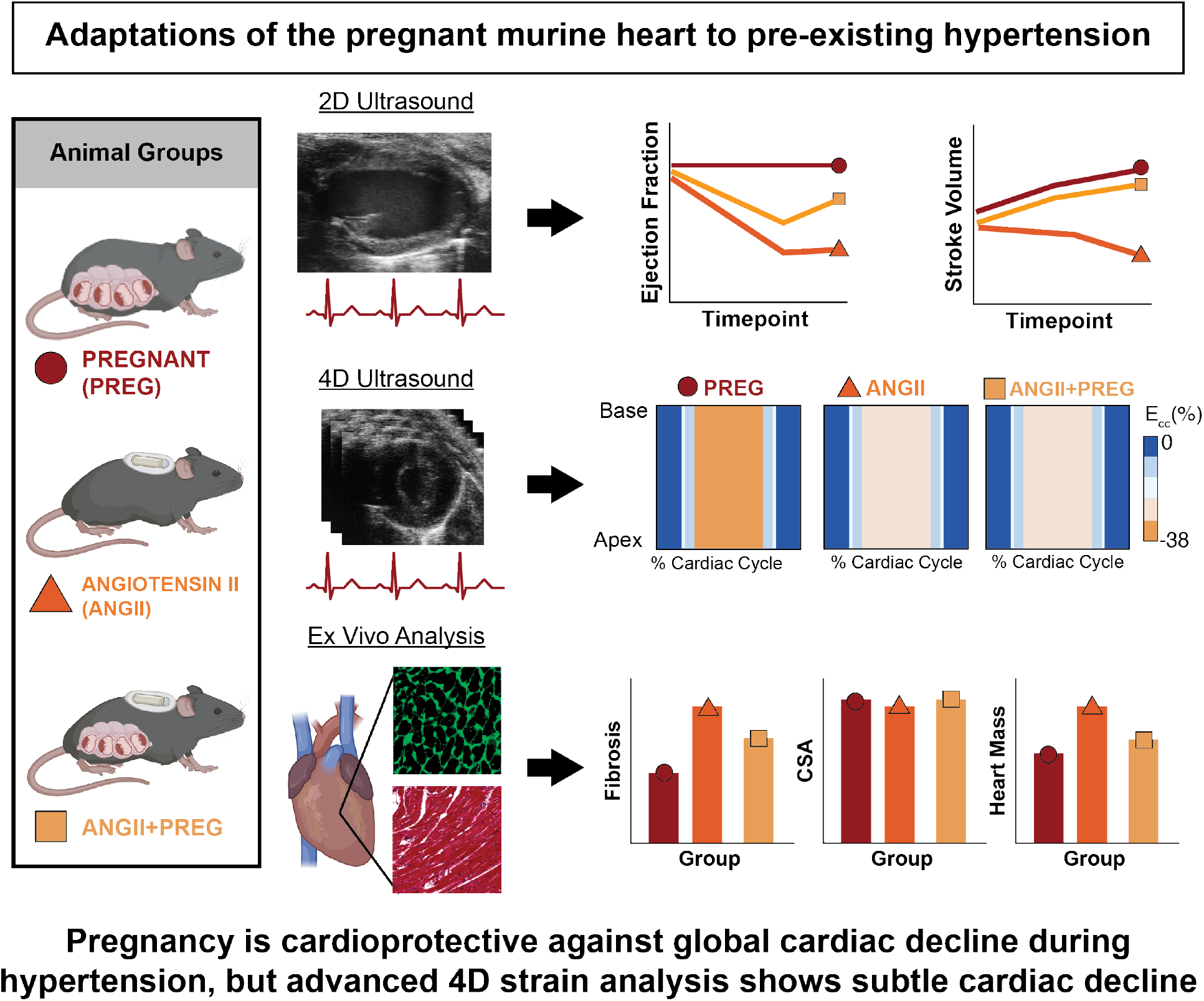

## 1 Introduction

Chronic hypertension occurs in 3-5% of all pregnancies [2, 1] and is associated with increased risk of developing preeclampsia, peripartum cardiomyopathy, and maternal and perinatal morbidity [2, 4]. Chronic hypertension during pregnancy is defined as pre-existing hypertension, hypertension that is diagnosed within the first 20-weeks of pregnancy, or hypertension that does not resolve within 12-weeks of delivery [5]. In healthy pregnancy, blood volume increases by approximately 50% and systemic vascular resistance decreases [6, 7]. The net effect of this volume overload is an increase in preload, or filling pressure, as well as an increase in stroke volume (SV) [8, 9]. In distinct contrast, hypertension is a form of pressure overload that increases resistance to left ventricle (LV) contraction (i.e., increased afterload), promotes myocardial fibrosis and thickening, and decreases SV [10, 11]. Interestingly, hypertensive disorders of pregnancy provide a unique environment in which pressure and volume overload co-exist and introduce competing effects on SV. Yet, the combined effect of these opposing stimuli on maternal cardiac health remains largely unknown [1].

Blood pressure naturally decreases in the latter half of the first trimester of pregnancy for both normotensive and hypertensive patients [2]. This attenuation and cardioprotective phenomenon is thought to be a downstream effect of vasodilation and reduced total peripheral resistance (TPR). Additional cardioprotective effects of pregnancy have also been observed in animal studies. For example, in a rat model of pregnancy-induced hypertension, cardiac collagen content was reduced in hypertensive pregnant rats compared to hypertensive controls [12, 2]. However, studies such as this exclude strain quantification, despite research showing that strain maladaptations in hypertensive patients may predate those of standard 2D metrics (e.g., ejection fraction) [13, 14]. Furthermore, the unique clinical differences [5] between chronic hypertension, gestational hypertension, early and late preeclampsia, and eclampsia are often overlooked [15]. Consequently, this leaves a gap in our understanding of the biomechanical and hemodynamic adaptations of the pregnant heart in response to chronic hypertension.

The objective of this study is to characterize the effects of hypertension on cardiovascular remodeling during pregnancy. To address these gaps, we quantified LV function, strain, and tissue morphology using *in vivo* 2D and 4D high-frequency ultrasound, and *ex vivo* histological analysis in a murine model of chronic hypertension during pregnancy. Results from this study provide critical insights into pregnancy-specific mechanisms of cardiac remodeling and may eventually inform clinical management of hypertensive pregnancies.

## 2 Methods

### 2.1 Ethical approval

All animal procedures were approved by the Purdue University Institutional Animal Care and Use Committee (Purdue IACUC 2002002016).

### 2.2 Animal Groups and Experimental Overview

Female C57Bl6/J mice (aged 7-11 weeks old, Jackson Laboratories, Bar Harbor, ME) were assigned to pregnant control (PREG, n=11), angiotensin II-treated control (ANGII, n=7), or ANGII+PREG (n=8) groups. A subset of PREG animals (n=5) were used only for blood pressure and ultrasound image collection. Two of the ANGII+PREG dams did not become pregnant soon enough to accommodate the 42-day lifespan of the mini-osmotic pump and one ANGII+PREG dam died after pump implantation, prior to pairing. These three animals were therefore excluded from the study, reducing the ANGII+PREG group size to n=5. The experimental timeline is summarized in **Figure 1** and experimental procedures are detailed below.

**Figure 1:**
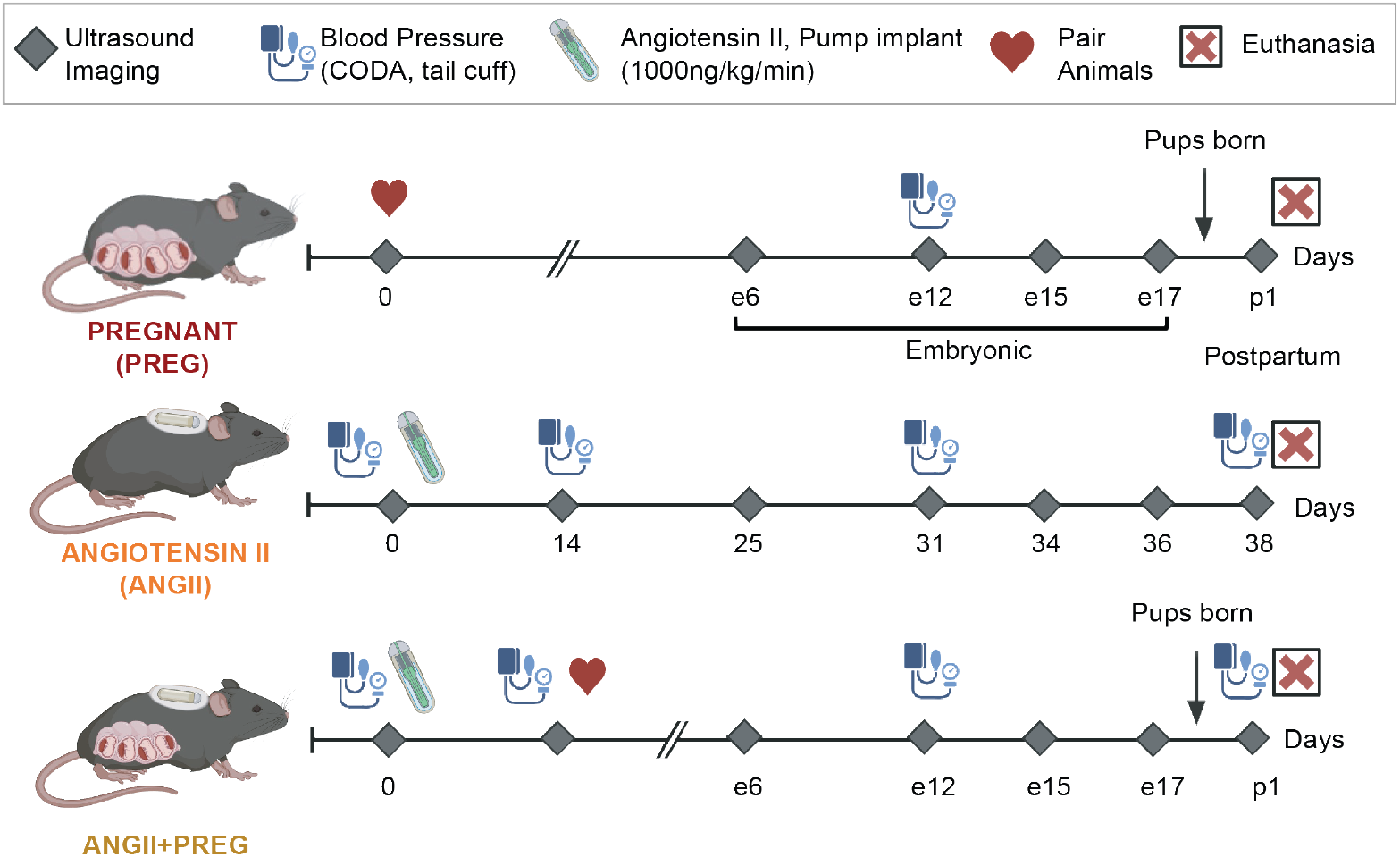
Experimental Timeline. Pregnant (PREG), angiotensin II treated (ANGII), and ANGII+PREG animals underwent ultrasound imaging and tail-vein blood pressure acquisition at baseline, on select embryonic days (e), and on postpartum day (p) 1. Experimental time-points for ANGII animals were matched to the ANGII+PREG cohort.

**Figure 2:**
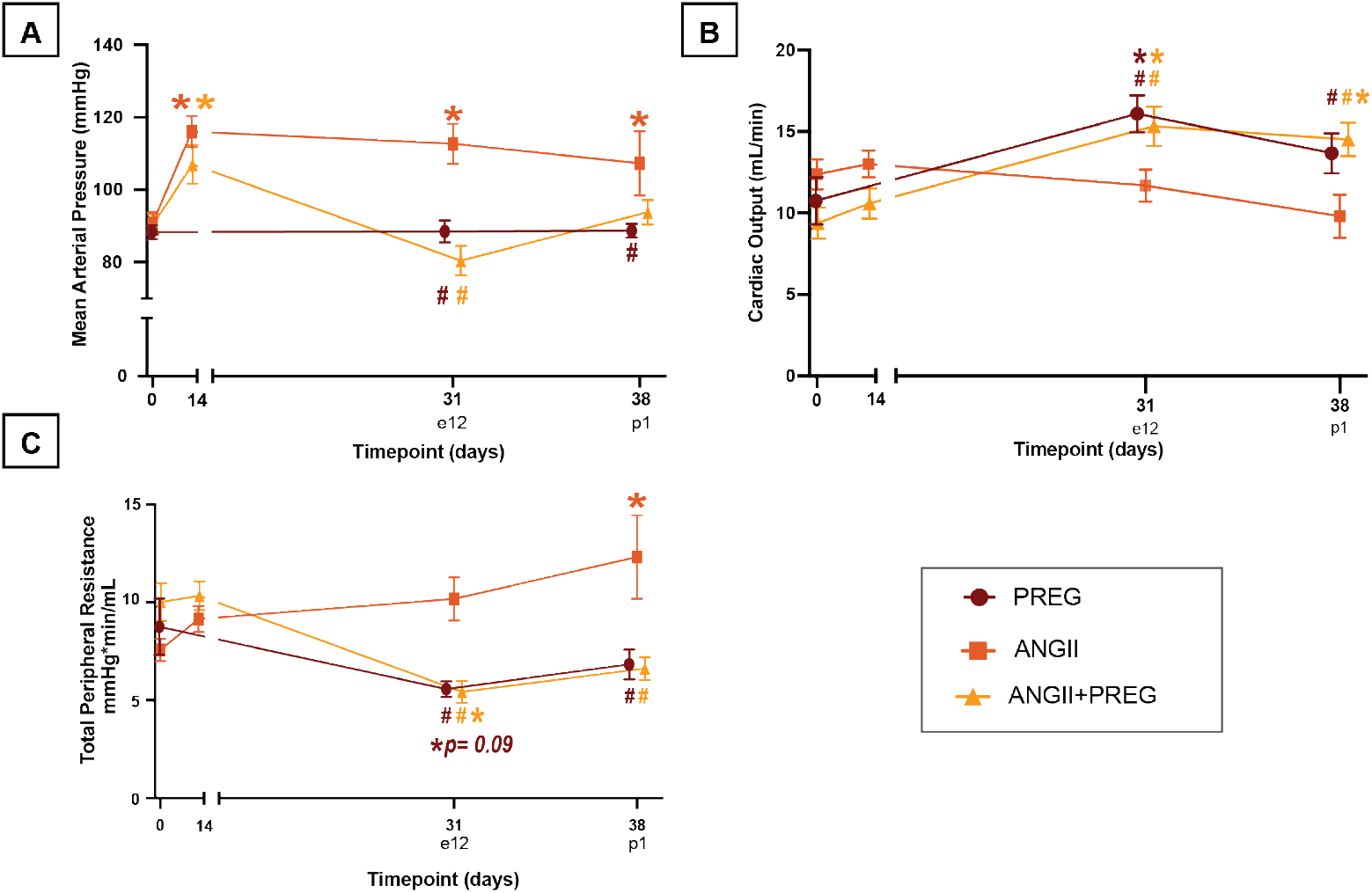
Changes in blood pressure and total peripheral resistance due to angiotensin II are modulated in late pregnancy. A) Mean arterial blood pressure (MAP), B) cardiac output (CO) measured from 2D ultrasound, and C) total peripheral resistance estimated from MAP and CO for PREG = pregnant control (n=5), ANGII = angiotensin II control (n=7), and ANGII+PREG= angiotensin II + pregnant group (n=5). \**p<*0.05 vs baseline, +*p<*0.05 vs pregnant cohort, #*p<*0.05 vs ANGII cohort. Color indicates group. Values=Mean*±*SEM.

### 2.3 Angiotensin II Infusion and Blood Pressure Acquisition

Mini-osmotic pumps (Model 2006, Alzet, Durect) were loaded with angII (1000ng/kg/min) and subcutaneously implanted in the dorsal space of the ANGII and ANGII+PREG animals for up to 42 days, as previously described [16]. Animals were anesthetized at 2-3% isoflurane during the procedure and given 0.05 mL of buprenorphine immediately after surgery. Blood pressure was measured via a tail-cuff system (2-channel CODA, Kent Scientific) in conscious animals at baseline prior to pump implantation, day 31/embryonic(e) day 12, and on day 38/postpartum(p) day 1. The ANGII and ANGII+PREG cohorts underwent additional blood pressure measurements 14 days after pump implantation to verify that hypertension was induced prior to continuing with the experimental procedure. Blood pressure was measured in the morning, prior to exposure to isoflurane, and within 24 hours of the associated imaging timepoint. Following previously established protocols [17], animals were placed in restraints on a heated stage and their tail temperature was maintained at 35-37°C. Animals were allowed to acclimate for 8-10 blood pressure measurement cycles. 10-20 blood pressure readings were collected per mouse and averaged at each experimental timepoint.

### 2.4 Breeding Scheme and Embryonic Dating

Female mice were housed together for 2-3 weeks to sync their estrous cycles prior to breeding. Dams in the PREG cohort were paired with male breeders after baseline imaging. Dams in the ANGII+PREG group were paired with male breeders 14 days after pump implantation once hyperension was established. All dams were paired with male C57Bl6/J mice in a 2:1 (female:male) breeding scheme. We performed abdominal ultrasound (MX550D, 25-55 MHz frequency bandwidth, Vevo 3100, FUJIFILM VisualSonics) of the uterine horns every 3-4 days to check for a decidual reaction and determine the embryonic day [18]. Dams were separated and moved to single housing after confirmation of pregnancy.

### 2.5 Image Acquisition and Analysis

All animals underwent serial imaging at: baseline prior to pump implantation and pairing, embryonic (e) days 12, 15, 17, and postpartum (p) day 1. Imaging timepoints for the ANGII control animals were matched to the ANGII+PREG animals based on the average time to conception. The average equivalent mini-osmotic pump day associated with the embryonic and postpartum imaging days were as follows: 25/e6, 31/e12, 34/e15, 36/e17, 38/p1. ANGII and ANGII+PREG animals also underwent image acquisition on day 14.

Animals were anesthetized with 1-3% isoflurane and placed in a supine position on a heat modulated imaging stage. Hair was removed using depilatory cream and heated ultrasound gel was applied to the imaging area. We collected long-axis (LAX) electrocardiogram-gated kilohertz visualization (EKV), and short-axis (SAX) 3D and 4D (3D+time) ultrasound images of the LV at each imaging timepoint. Image collection was performed within 24 hours of the scheduled imaging day. Temperature, respiratory rate, and heart rate were monitored while imaging.

Ejection fraction (EF), end diastolic volume (EDV), peak systolic volume (PSV), stroke volume (SV), and cardiac output (CO) were measured from 2D EKVs using VevoLab (Version 5.7.1, FUJIFILM VisualSonics). These values were measured based on end diastolic and peak systolic endocardial tracings of the LV. Circumferential (*E_cc_*) and longitudinal (*E_ll_*) strain, and strain rates (SR) were quantified from 4D images using a custom MATLAB graphical user interface [19]. In brief, the endocardial and epicardial borders of the LV were segmented from base to apex across the cardiac cycle (6 rotations per slice, 4 slices from base to apex). A volume mesh was created from these 48 points and used to calculate strain across the cardiac cycle. Global strains were measured as the average strain across the LV. Strain rates were measured as the slope of the strain-time curve, the rate of change in strain [20, 21].

### 2.6 Litter Size, Tissue Collection, and Histology

We recorded litter size and pup weight at the time of euthanasia (day38/p1). Dams were euthanized with an overdose of isoflurane followed by a thoracotomy. The inferior vena cava was then perfused with 0.1 mg/mL of potassium chloride to arrest the heart in diastole. We excised and weighed whole hearts, followed by fixation in 4% paraformaldehyde for 48 hours. After initial fixation, hearts were transferred to 70% ethanol until they were processed for histology. Hearts were sectioned in 4-5 segments from base to the apex along the short axis, embedded in paraffin, and sliced. Tissue samples were stained using hematoxylin and eosin (H&E), Masson’s trichrome (MTC), and Wheat-Germ Agglutinin (WGA) staining and scanned at 20x magnification. H&E and WGA stained sections were evaluated by a board certified veterinary pathologist for qualitative assessment and quantitative fibrosis analysis, respectively. Percent fibrosis was measured across all MTC sections using the ImageScope color deconvolution tool (Leica Biosystems). Cardiomyocyte cross-sectional area (CSA) was quantified from WGA-stained samples. CSA analysis was performed in eight randomly selected regions of interest (ROI) along the endocardium and epicardium at mid-papillary. Within each ROI, CSAs were measured using the Analyze Particles function and averaged together [22]. A size range of 60*µ*m^2^-infinity and a circularity of 0.40-0.80 was utilized to exclude non-cardiomyocyte structures, including vasculature. CSA was not analyzed in two hearts from the PREG group and two hearts from the ANGII group due to inadequate staining quality.

### 2.7 Statistical Analysis

We performed statistical analysis in Prism 10 (GraphPad Software, San Diego, California, USA) with *p<*0.05 indicating significance. We performed mixed effects model analysis with post hoc Tukey’s multiple comparisons test to compare values across timepoints both within and across all animal groups. Values that were collected at a single timepoint, such as litter size and heart mass, were assessed with a one-way repeated measure analysis of variance (ANOVA). Non-parametric equivalents were used for non-normal data. All results are reported as mean*±*standard error of mean (SEM).

## 3 Results

### 3.1 Dam & Litter Outcomes

Although animals were similar in age, body mass was higher at baseline for the ANGII group (19.29 *±* 1.11g) compared to the PREG (17.38*±*0.52g, *p*=0.04) and ANGII+pregnant (16.92 *±* 0.64g, *p*=0.04) group (**Supplemental Fig. S1A**). Despite this initial difference, body mass was similar between all three groups on day 38/p1 at approximately 22g. Both the pregnant and ANGII+PREG groups experienced significant increases in body mass by day 25/e6 compared to their baseline (PREG: 19.87*±*0.29g, *p*=0.035; ANGII+PREG: 21.64*±*0.46g, *p*=0.001) and weighed more than the ANGII group from day 31/e12 up until the final pregnancy imaging timepoint (day 36/e17). Body mass was similar throughout pregnancy for both the PREG and ANGII+PREG groups, peaking at 30.92*±*0.51g for pregnant and 31.90*±*0.94g for ANGII+PREG animals on day 36 (e17). No significant differences in mean pup weight were observed between the PREG and ANGII pregnant groups (PREG: 1.17*±*0.01g, PREG+ANGII: 1.24*±*0.06g) or in litter size (PREG: 6.73*±*0.47 pups, PREG+ANGII: 5.80*±*1.11 pups) **(Supplemental Fig. S1B&C)**.

### 3.2 Blood pressure and total peripheral resistance are subdued in ANGII+PREG

Mean arterial pressure (MAP) was maintained at approximately 88.5*±*2.8mmHg in the PREG group across the experimental timeline. Conversely, in the ANGII group, MAP increased to 112.7*±*5.6mmHg by day 31 (*p*<0.001) and slightly decreased at day 38 (107.2*±*8.9mmHg, *p*=0.01) compared to baseline. In ANGII+PREG animals, despite the initial increase in blood pressure due to angII **(Supplemental Fig. S2A-C)**, blood pressure normalized at day 31/e12 and matched MAP measured in the PREG group (88.6*±*1.9mmHg). Systolic and diastolic blood pressures are available in **Supplemental Fig. S2 and Table S1**.

We estimated TPR from tail-cuff-based MAP and CO measured from 2D LAX EKV images. In the PREG group, TPR mildly decreased by day 31/e12 (5.6*±*0.4mmHG*min/mL, *p*= 0.09). In ANGII, TPR increased by day 38/p1 compared to baseline (12.3*±* 2.1mmHG*min/mL, *p*=0.003). TPR significantly decreased in the ANGII+PREG group by day 31/e12 (5.6*±*0.4mmHG*min/mL, *p*=0.018) and matched the negative trend observed in the PREG group. Although the changes for PREG and ANGII were not significant compared to their respective baselines, TPR was significantly lower in both the PREG (*p*=0.013) and ANGII+PREG (*p*=0.009) compared to ANGII animals at day 31/e12.

### 3.3 Cardiac function partially adapts in ANGII+PREG

LV volumes and EF were measured from 2D LAX EKV images by tracking the endocardial wall at end diastole and peak systole (**Fig. 3A**). PSV remained stable in PREG animals with no significant changes relative to baseline (**Fig. 3C**). PSV increased in both the ANGII and ANGII+PREG groups and approximately peaked on day 31/e12(ANGII: 24.67*±*3.11*µ*L, *p*=0.118; ANII+PREG: 29.82*±*3.31*µ*L, *p*=0.032). PSV was significantly greater in the ANGII (22.20*±*2.76*µ*L, *p*=0.002) and ANGII+PREG (20.64*±*2.18*µ*L, *p*=0.018)groups compared to PREG animals (10.59*±*0.65*µ*L) as early as day 25/e6. EDV increased in all groups relative to baseline (Fig. 3B). EDV increased compared to baseline until day 31/e12 before plateauing in the PREG group (44.60*±*1.54*µ*L, *p*=0.019) and ANGII group (46.53*±*2.33*µ*L, *p*=0.028). In contrast, the ANGII+PREG group showed more sustained increases until day 38/p1 (50.87*±*2.12*µ*L, *p*=0.023) and was significantly higher than PREG animals by day 34/p1. SV increased from baseline to postpartum by 6.2*µ*L in PREG (*p*=0.371) animals and by 13.8*µ*L in ANGII+PREG (*p*=0.041) animals (**Fig. 3D**). Conversely, SV decreased by 5.21*µ*L in ANGII animals throughout the experimental timeline, though not significantly (*p*=0.104). EF did not change in the PREG group and was maintained at approximately 65%. EF decreased in the ANGII group compared to baseline and was significantly lower than in the pregnant group by day 25/e6. EF was also significantly lower in ANGII+PREG animals compared to PREG animals until day 31/e12. However, this decrease was not sustained. At day 34/e15, mid-pregnancy, EF began to increase in the ANGII+PREG group and trended towards PREG measured values. LV volumes and EF are reported in Supplemental Table S2.

**Figure 3:**
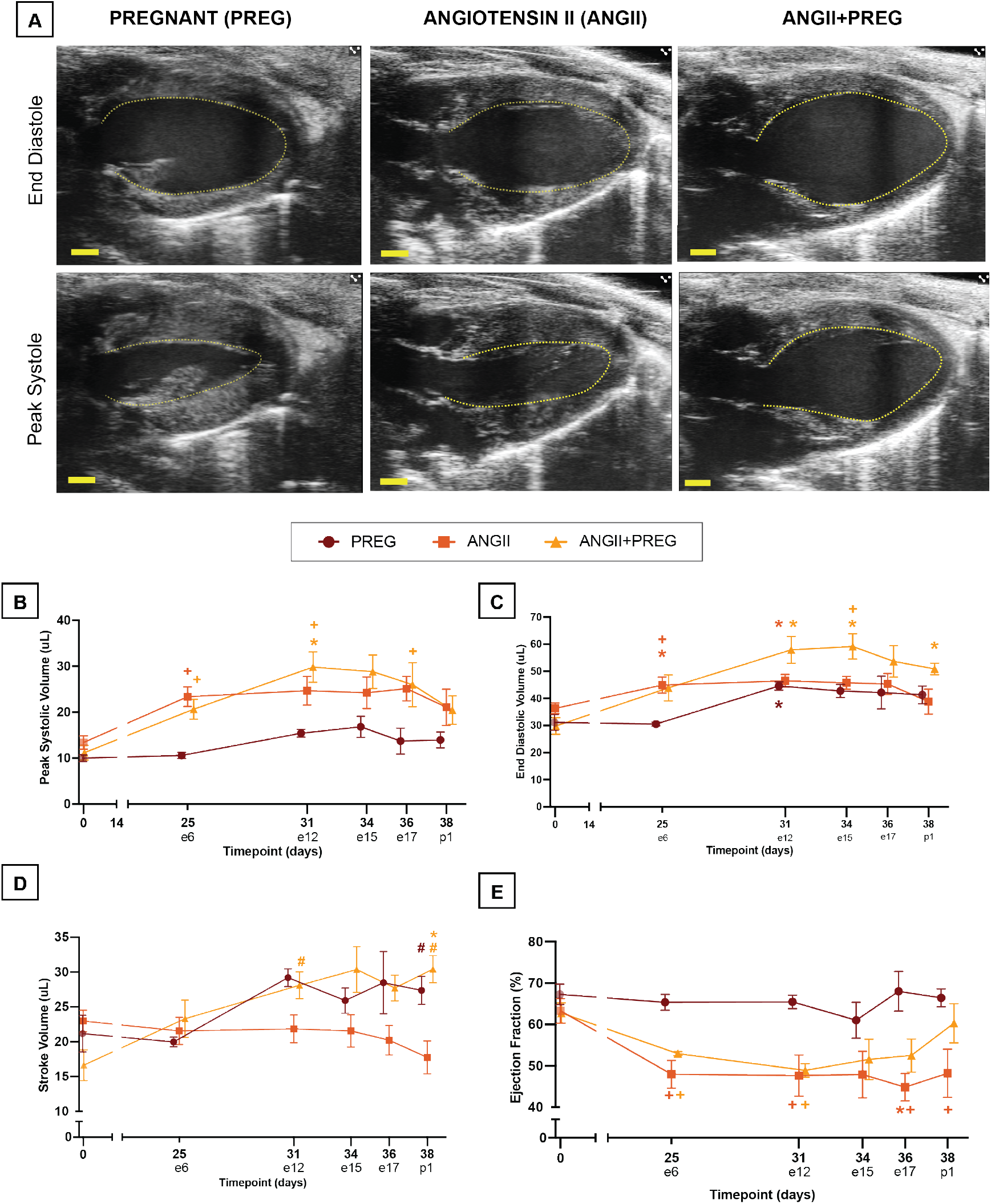
Stroke volume and ejection fraction adapt to angiotensin II during pregnancy. A) Representative 2D ECG-gated kilohertz visualization images of the left ventricle at end diastole and peak systole. Scale bar= 1mm. Endocardium is outlined in yellow-dotted line. B) End diastolic volume. C) Peak systolic volume. D) Stroke volume measured from EDV and PSV. E) Ejection fraction measured from SV and heart rate. PREG= pregnant control (n=4-5), ANGII= angiotensin II control (n=7), ANGII+PREG= angiotensin II+pregnant group(n=4-5). \**p<*0.05 vs. baseline, +*p<*0.05 vs. pregnant cohort. Color indicates group. Values=Mean*±*SEM.

### 3.4 Left ventricle strain mirrors peak systolic volume

We first visualized strain with Cartesian heatmaps of *E_cc_* and *E_ll_* across the entire the LV (y-axis) with respect to the cardiac cycle (x-axis) (**Fig. 4**). From these, we observed a homogeneous strain distribution across the ventricle within each group (**Fig. 4 A&B**). Heatmaps of the ANGII and ANGII+PREG group showed a visible decrease in maximum strain (lighter orange and white regions) during systole, reflective of reduced left ventricular contraction. Raw heatmaps for *E_cc_* and *E_ll_* are available in **Supplemental Fig. S3-S8.** We quantified the kinematic behavior of the ventricle observed in heatmaps by measuring global strains (**Fig. 4C&D, Table S3**). In PREG animals, global *E_cc_* and *E_ll_* were maintained at approximately -31% and -22%, respectively. In contrast, global *E_cc_* significantly decreased from baseline for ANGII (-24.22*±*1.48%, *p*=0.048) and ANGII+PREG (-26.94*±*1.65%, *p*=0.044) on day 25/e6. In late pregnancy, ANGII (-22.0*±*2.42%, *p*=0.019) and ANGII+PREG (- 23.43*±*0.76%, *p*=0.003)animals showed significantly lower, global *E_cc_* compared to PREG animals. Global *E_ll_* was also lower in ANGII (-15.83*±*1.13%, *p*=0.002) and ANGII+PREG (-14.48*±*0.81%, *p*=0.001) animals in late pregnancy (e17/day36). Similar trends were observed in regional strains (**Supplemental Fig. S9**).

**Figure 4:**
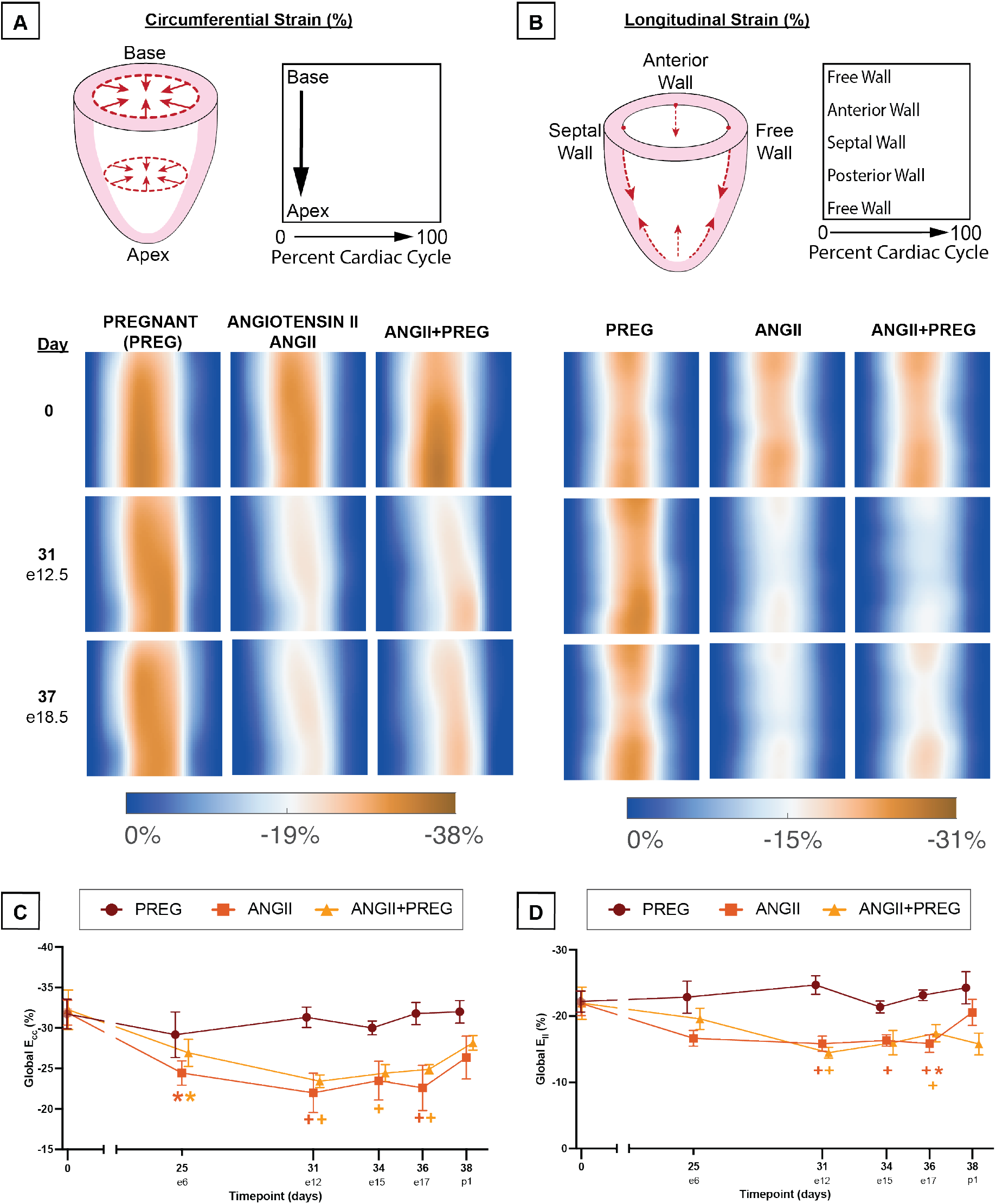
Left ventricle strain decreases during angiotenisn II combined with pregnancy. A-B) Schematic of circumferential- and longitudinal strain and average heatmaps at embryonic (e) day 12.5 and 15.5. C) Global circumferential- and longitudinal strain across experimental timeline. PREG= pregnant control (n=4-5), ANGII= angiotensin II control (n=7), ANGII+PREG= angiotensin II+pregnant group (n=4-5). \**p<*0.05 vs. baseline, +*p<*0.05 vs. pregnant cohort. Color indicates group. Values=Mean*±*SEM.

### 3.5 Early diastolic strain rate decreases with angII infusion

To assess the rate of contraction and relaxation, we quantified systolic strain rate (sSR) and early diastolic strain rate (dSR) (**Fig. 5, Supplemental Table S3**). Within each group, sSR was primarily preserved compared to baseline (**Fig. 5B&C**). However, sSR was significantly lower in ANGII and ANGII+PREG cohorts when compared to the PREG group starting on day 31/e12. In PREG animals *E_cc_* sSR was -1.57*±*0.09% on e12/day31, while the ANGII and ANGII+PREG cohorts were significantly lower at -1.06*±*0.15% (*p*=0.036) and -0.91*±*0.03% (*p*=0.003), respectively. For *E_ll_* sSR on day12/day31, PREG was -1.10*±*0.07%, while ANGII and ANGII+PREG cohorts were significantly lower at -0.74*±*0.07% (*p*=0.009) and -0.65*±*0.03%, (*p*=0.003), respectively. *E_cc_* dSR increased in PREG animals compared to baseline, peaking on day 31/e12 (2.26*±*0.12%, *p*=0.006)(**Fig. 5D&E**). ANGII+PREG animals did not share this increase in *E_cc_* early dSR. On day 34/e15 *E_cc_* dSR was significantly lower in ANGII animals (1.12*±*0.06%) compared to PREG animals (1.97*±*0.10%, *p*<0.001). ANGII and ANGII+PREG animals saw significant mid-gestational (day 31/e12) decreases in *E_ll_* dSR versus baseline (*p*=0.020 and *p*=0.012, respectively), while the PREG cohort saw significant increases. This divergence of PREG vs ANGII and ANGII+PREG animals was statistically different on on day 31/e12 and day 34/e15.

**Figure 5:**
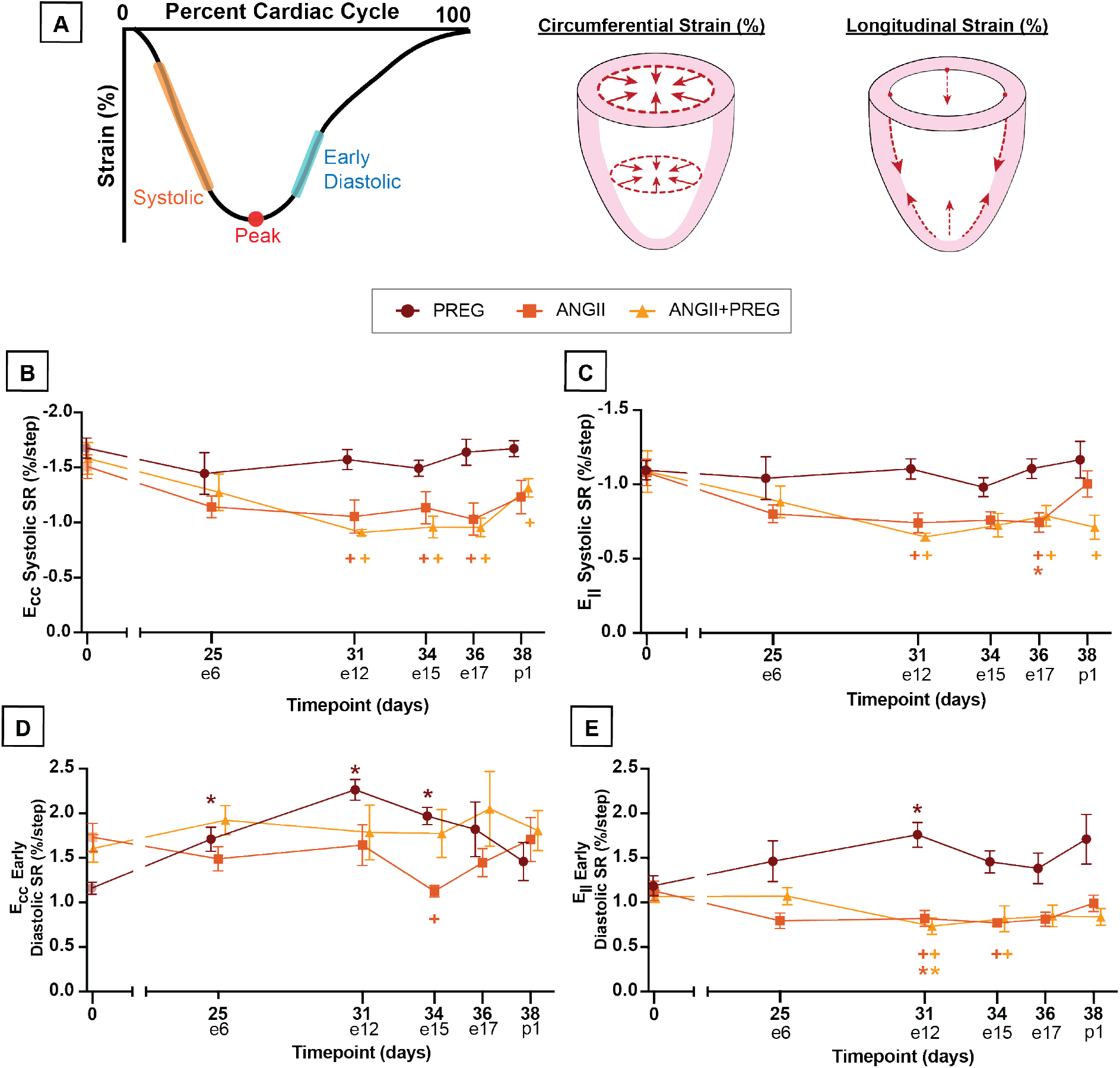
Strain rate decreases with exposure to angiotensin II. A) Schematic of strain rates on representative strain-time curve. B-C) Systolic and D-E) early diastolic strain rate (SR) for circumferential (*E_cc_*) and longitudinal (*E_ll_*) strain. PREG= pregnant control (n=4-5), ANGII= angiotensin II control (n=7), ANGII+PREG= angiotensin II+pregnant group (n=4-5). Values=Mean*±*SEM.

### 3.6 Angiotensin II induced minimal fibrotic remodeling

Representative histological images at 2x and 20x for PREG, ANGII, and ANGII+PREG can be seen in **Fig. 6A-C**. One out of seven ANGII animals and two out of five ANGII+PREG animals had myocardial lesions, consisting of multifocal areas of myocardial fiber disarray with varying myocardial fiber size and orientation (**Fig. 6D**). MTC stains of these mural myocardial lesions confirmed minimal amounts of fibrosis in the area. We observed changes to the myocardial arteries of one animal in the ANGII group and one animal in the ANGII+PREG group. Myocardial arteries of these mice were multifocally affected and had increased perivascular cells, smooth muscle degeneration, subendothelial edema, and prominent endothelial cells. The MTC stain of these arterial lesions indicated that minimal fibrosis surrounds the affected arteries. No lesions were identified in the PREG hearts stained with H&E and MTC. Quantitative analysis of fibrosis across select ROIs showed significant differences between PREG and ANGII animals (2.9*±*0.1% vs. 5.6*±*0.6%, *p*=0.011) (**Fig. 6E**). The percent fibrosis was lower in ANGII+PREG (4.8*±*0.6%) animals compared to ANGII alone and higher than in PREG animals, though not significantly. It is important to note that the biological relevance of the quantitative fibrosis scores may be limited due to the lack of qualitative findings. Lesion presentation and fibrosis percentages are reported in Supplemental Table S4.

**Figure 6:**
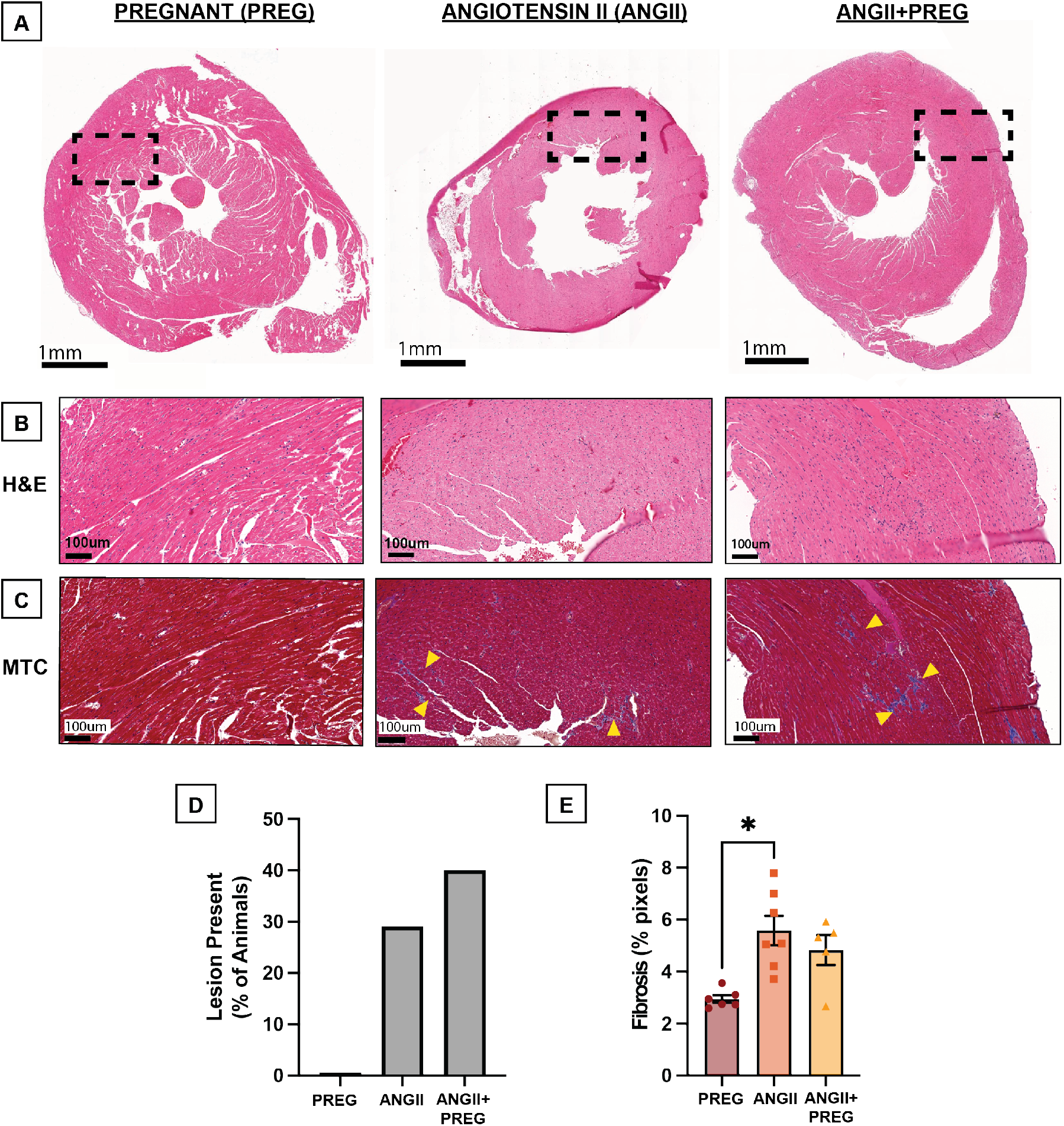
Fibrosis is minimal in angiotensin II (ANGII) and ANGII+Pregnant animals. Left ventricle histology of A) with Hematoxylin and Eosin (H&E) at 2X. Dotted line indicates region shown at 20x in B-C) for H&E and Masson’s Trichrome. Yellow triangles indicate regions with higher collagen content. D) Presence of lesions at excision. E) Quantitative fibrosis percentage of each excised heart. n=6 for pregnant controls (PREG), n=7 for angiotensin II control (ANGII), and n=5 for ANGII+PREG group. Values=Mean*±*SEM.

### 3.7 Cross-sectional area was unchanged and heart mass was altered

Representative WGA images used for CSA analysis, including outlined ROIs, are shown in **Figure 7A&B**. Average CSA was similar in ANGII (188.0*±*6.8 *µ*m^2^) and ANGII+PREG animals (188.0*±*8.0 *µ*m^2^) **(Fig. 7C)**. The PREG group had a larger average CSA of 199.6*±*8.9 (*µ*m^2^)); however, this was not significantly different from either ANGII or ANGII+PREG animals. CSAs are reported in Supplemental Table S4. Raw excised heart mass was significantly greater in the ANGII group (190.67*±*9.44mg) compared to both the PREG (137.47*±*6.20mg, *p<*0.001) and ANGII+PREG (157.98*±*4.06mg, *p*=0.015) groups **(Fig. 7D)**. No significant differences were noted in heart mass between PREG and ANGII+PREG animals. This observation held true even when normalized to body mass measured on the day of tissue collection.

**Figure 7:**
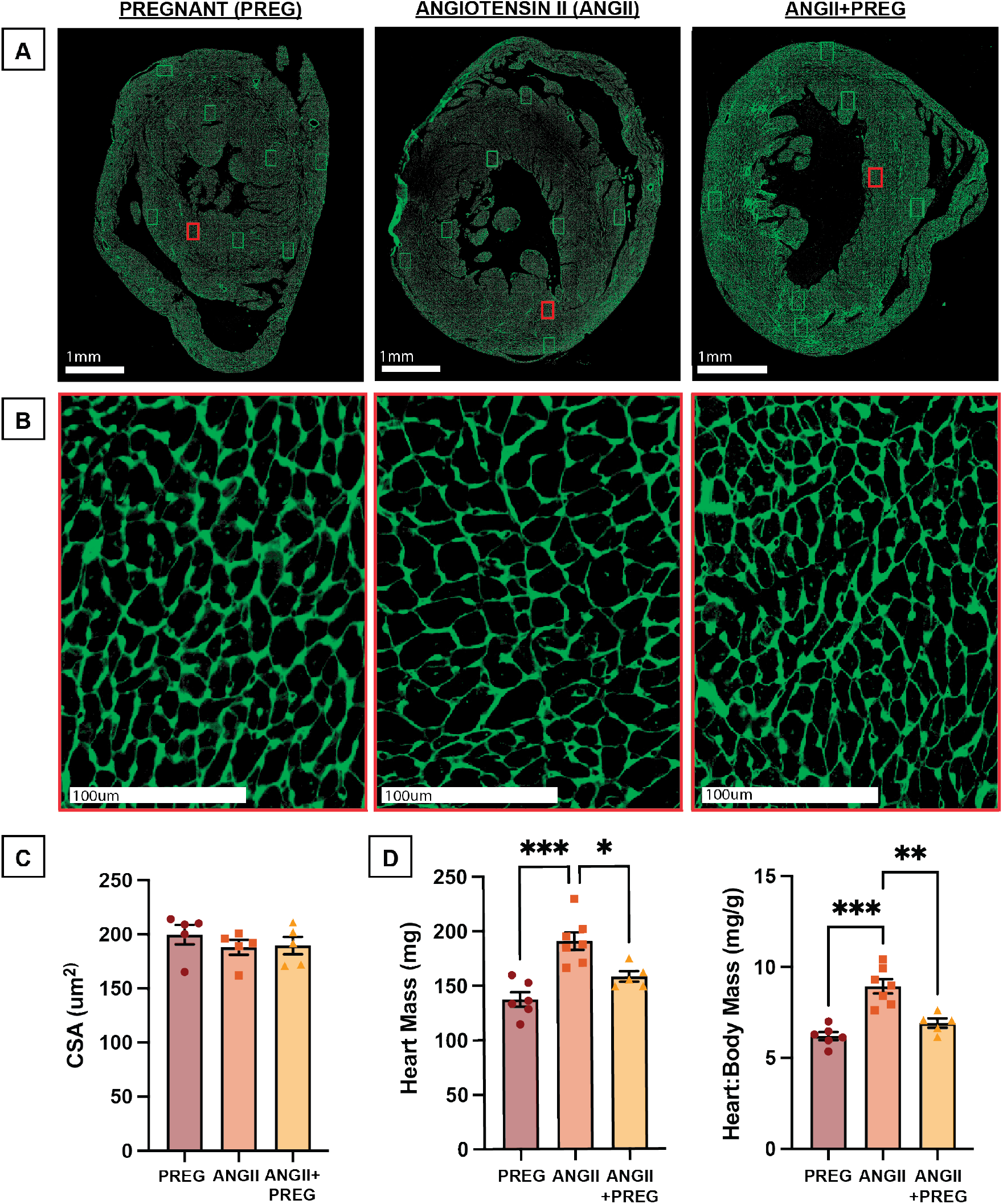
Cardiac hypertrophy from angiotensin II is subdued during pregnancy. Wheat-Germ Agglutinin stains of representative short-axis left ventricle slices at mid-papillary at A) 2X with analyzed regions of interest outlined in boxes and at B) 20X corresponding to the red rectangles. C) Cardiomyocyte cross-sectional area, n=5/group. D) Heart mass and normalized heart mass, n=5 Values=Mean*±*SEM.

## 4 Discussion

In this study we investigated the effects of combined pressure and volume overload in hypertensive murine pregnancy on cardiac function, strain, and morphology. From excised heart mass, tail cuff blood pressure, *in vivo* high-frequency ultrasound imaging, and histological analysis, we observed an adaptive remodeling response in angII-infusion superimposed with pregnancy. In the ANGII+PREG group, we observed pregnancy-related protective cardiac adaptations through SV and heart mass, but observed reductions in cardiac strain and strain rate with greater variability between animals. The cardiac changes described here provide novel insights into the potential protective response of the pregnant heart to chronic hypertension and emphasize the importance of using a multi-modal approach to assess cardiac health during pregnancy.

### 4.1 Blood pressure and TPR adapt during angII-exposed pregnancy

Blood pressure and TPR showed positive adaptations during pregnancy that overcame the effects of angII. Reflecting the vasoconstriction and sodium and water retention induced by chronic angII infusion [23, 24], MAP significantly increased in ANGII and ANGII+PREG animals after 14 days of angII infusion. However, the observed increases in MAP were mild relative to the clinical definition of hypertension (systolic blood pressure *≥*140mmHg, diastolic blood pressure *≥*90mmHg [25, 26]). Despite continued exposure to angII, MAP normalized in ANGII+PREG animals by mid-pregnancy. These results are consistent with clinical studies [27, 28] that show a natural decrease in blood pressure during mid-pregnancy in both normotensive and hypertensive individuals, and in rodent data [29, 30, 31]. It is believed that mid-gestational decreases in BP may be due to increased levels of circulating progesterone and estrogen during pregnancy, which decrease sensitivity to angII and reduce systemic vascular resistance [32, 33, 34]. This may explain the protective effects that we observed in pregnancy against angII infusion.

In humans, the first trimester TPR drop is thought to initiate compensatory changes of increased CO and blood volume during pregnancy [35]. In line with this, TPR decreased in PREG and recovered in ANGII+PREG animals. In a renal artery ligation model comparing short-standing and established hypertension during pregnancy, Lundgen *et al.*, similarly observed decreased TPR in hypertensive-pregnant animals compared to normotensive controls. [36]. However, unlike the rats with short-standing hypertension, we did not see higher TPR compared to normotensive pregnancy. This may be due to more severe elevations in MAP in the ligation model compared to our angII model. Clinically, TPR is also elevated in hypertensive patients compared to normotensive controls [37], but unlike the animals models, TPR does not decrease mid-gestation in hypertensive patients [37, 15]. The TPR recovery seen in both animal models, unlike in clinical data, could indicate that murine models may have different biochemical adaptations in pregnancy compared to humans. Notably, TPR has been shown to be a valuable predictor of complications later in pregnancy, such as preeclampsia [38, 39, 40]. In this context, our findings may suggest TPR assessment during pregnancy could help elucidate differences in the dynamic maternal vascular response.

### 4.2 Global cardiac function is attenuated in late pregnancy despite angII infusion

Maternal blood volume increases by nearly 50% during gestation [41] and subsequently causes an increase in diastolic filling pressure. Reflecting this physiological adaptation and consistent with Kaissar *et al.* [42], EDV increased in PREG animals relative to their baseline. EDV has also been shown to increase in male mice treated with angII when measured with cardiac magnetic resonance imaging [43], which aligns with our ANGII animals. Notably, EDV showed the most substantial increase in ANGII+PREG animals, possibly due to combined pressure and volume overload. We observed a mild increase in PSV in both ANGII and ANGII+PREG animals, which may reflect slightly reduced systolic function, as similarly observed with LV diameter [44]. As the ventricle loses the ability to fully contract during systole (i.e, reduced wall motion), more blood is left in the ventricle after peak contraction. Pressor doses of angII infusion in rodents have been shown to reduce CO, EF, and SV [45, 43, 46]. Interestingly, SV recovered in late pregnancy in ANGII+PREG animals, suggesting that global cardiac function can positively adapt in pregnant animals with chronic angII exposure. These results differ from Morgan, *et al.,* who reported decreased SV in pregnant stroke-prone spontaneously hypertensive (SHRSP) rats treated with angII [45]. This observed difference in the functional response of the heart could be attributed to the use of SHRSP rats, which have chronically elevated blood pressure. The heart’s ability to adapt to elevations in blood pressure may therefore depend on the severity of the hypertensive phenotype. Taken together, our observations from 2D ultrasound suggest that SV can overcome angII related decreases during pregnancy, but that more detailed cardiac measures such as PSV and EDV may indicate early maladaptive remodeling.

### 4.3 Strain analysis shows subtle cardiac decline

Our animal model provides several valuable advantages. Using *in vivo* ultrasound, we were able to collect serial, longitudinal data that captures not only LV function, but also strain mechanics. This is the first report of 4DUS-derived strain changes in a model of pre-existing hypertension during pregnancy. While traditional global measures of cardiac health, such as EF, suggest that the pregnant heart can overcome adverse functional changes due to angII, our advanced 4D strain measures revealed adverse LV remodeling. Notably, global *E_cc_*, *E_ll_*, and sSR were lower in ANGII and ANGII+PREG groups compared to pregnancy alone and could be attributed to the observed increases in PSV. Previous literature reports modest decreases in global strain in late normotensive pregnancy that rebounds during postpartum [47, 48]. Our data is in agreement with both rat and clinical studies showing significantly reduced global *E_ll_* in preeclamptic patients and patients experiencing hypertensive disorders of pregnancy compared to normotensive pregnancies [49, 50, 51, 52, 53]. Although strain adaptations were small compared to baseline, these findings remain noteworthy as strain was able to provide both directional and regional information in all groups throughout pregnancy. Our 4D analysis also revealed changes in diastolic function, as measured by dSR. Pregnancy caused an increase in dSR, similar to volume overload induced with an IVC-to-aortic shunt [54]. Conversely, our dSR was lower in our ANGII group. This is in agreement with murine pressure overload induced via abdominal aortic banding, which has been shown to cause a decrease in longitudinal dSR [54]. This decrease in dSR may suggest impaired LV isovolumetric relaxation, and thus, reduced ventricle filling. Interestingly, in ANGII+PREG animals, decreases in dSR were not accompanied by the expected decrease in EDV. This dichotomy suggests that dSR may be a more sensitive measure of diastolic function compared to EDV, particularly in cases where volume overload overshadows decreases in ventricle filling volume caused by pressure overload. Overall, these results suggest that 2D cardiac ultrasound and blood pressure measurements may not fully capture cardiac remodeling and that more nuanced functional measures should be considered when assessing maternal cardiac health.

### 4.4 Angiotensin II leads to minimal cardiac fibrosis

It is well established that chronic hypertension can lead to a pathological cardiac hypertrophic response with irreversible fibrotic deposition [55]. Additionally, proposed theories suggest there are competing pathologic and physiologic cardiac hypertrophy signaling pathways in hypertensive pregnancies [56]. Specifically, hormonal (estrogen and progesterone) and PI3K/Akt signaling may attenuate the angII/AT1R pathways that increase vasoconstriction [56]. In line with this, we measured an increase in cardiac fibrosis in ANGII animals compared to PREG animals, and a slight reduction in ANGII+PREG animals. These results are supported by a similar study by Aljabri *et al.,* in which rats infused with angII (150 ng/kg/min) on day 6 of gestation showed a dramatic decrease in cardiac collagen compared to hypertensive controls when observed via Sirius Red staining [12]. It should be noted, however, that the minimally visible collagen deposits in our histological studies suggest limited pathological significance. Therefore, care should be taken to consider the overall severity of fibrosis and to not rely on quantitative analysis alone. Additionally, Aljabri *et al.* observed a more pronounced difference between angII-treated pregnant rats compared to both pregnant and angII-treated controls. This difference between our mice and Aljabri’s study, may be due to higher systolic and diastolic pressures that were observed in the rat-based study. Furthermore, wall stress is directly proportional to radius 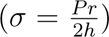, thus it is conceivable that the smaller LV radius in mice compared to rats may not be as capable of stressing the cardiomyocytes at the same angII dose.

### 4.5 Pregnancy attenuates adverse changes to cardiac hypertrophy during exposure to angiotensin II

Increased LV mass has been shown to occur during normotensive pregnancy [57, 58] as the heart accommodates increased preload, i.e., myocardial stretching during diastole. Moreover, angII treatment in mice has also been shown to cause an increase in cardiac mass [59]. However, the effects of combined pregnancy and angII infusion did not compound the degree of cardiac growth. When angII was combined with pregnancy, cardiac mass was lower compared to non-pregnant angII-treated mice and was similar that of normotensive pregnant mice, suggesting that hypertension-induced cardiac hypertrophy is reduced in settings of pregnancy. The attenuation in blood pressure during pregnancy, and therefore less severe pressure overload, may have contributed to lower cardiac mass in ANGII+PREG animals compared to their non-pregnant counterparts. Cardiomyocyte analysis from these same hearts showed no difference in the average CSA. This matches the results of the Aljabri *et al.* rat study, which reported no differences in cardiomyocyte diameter between sham, pregnant, ANGII, and ANGII+pregnant rats [12]. One explanation for the inconsistent findings between heart mass and CSA could be that the eccentric hypertrophy seen in pregnancy and pressure overload can lead to increased heart mass with little cardiomyocyte thickening [60, 61].

### 4.6 Limitations and Future Work

An overarching limitation of our work is the challenge of aligning the timeline of the miniosmotic pump to the embryonic days of pregnancy. To encourage timely conception, we housed dams together prior to breeding to synchronize their estrous cycles, provided mouse huts, added soiled bedding from male breeders, and minimized cage disruptions. Despite these precautions, it is not possible to precisely control the time from pairing to conception. This resulted in slight variability in the number of days the mini-osmotic pump was present before the onset of pregnancy in the ANGII+PREG cohort. To standardize timelines between the PREG and ANGII+PREG groups, we scheduled data collection according to embryonic stages confirmed via uterine ultrasound. However, since ANGII animals were not pregnant, corresponding embryonic timepoints did not exist. To address this, we calculated the average number of days between pairing and conception in the ANGII+PREG group and used this to calculate an average timeline relative to mini-osmotic pump implantation that was used as a matched timeline for ANGII animals. Blood pressure increased significantly with the introduction of angII, but these increases were not sustained and did not meet the definition of hypertension (systolic blood pressure *≥*140mmHg, diastolic blood pressure *≥*90mmHg [25, 26]). Future work could utilize a transgenic rodent strain to better mimic chronic hypertension and to induce more severe increases in blood pressure [62]. This would additionally eliminate the limitation of the 42-day lifespan of the mini-osmotic pumps, and consequently, allow the experimental timeline to be extended into the the postpartum period. Additionally, although hearts were perfused, excised heart mass may be affected by remnant blood in the ventricle and is less sensitive than CSA analysis. Within our experimental timeline, pregnancy partially preserved cardiac function under pressure overload. Future studies could assess whether compensatory mechanisms continue in postpartum relative to lactation. Other hypertensive disorders during pregnancy, such as preeclampsia, have distinct clinical definitions and physiological effects [5]. These differences may lead to cardiovascular adaptations that differ from those observed in our animal model of hypertension prior to pregnancy.

## 5 Conclusion

In summary, we measured cardiac function, strain, and blood pressure in a longitudinal study of pregnancy, chronic hypertension, and pregnancy complicated by pre-existing hypertension. Our ultrasound-based analysis suggests that pregnancy alters cardiac function and remodeling in response to increases in blood pressure. Although LV strain decreased in animals with angII exposure during pregnancy, hypertrophy decreased, and both blood pressure and left ventricular function began to normalize by mid-gestation. Thus, the interplay of pregnancy and angII creates a unique environment of combined pressure and volume overload in which the maternal heart adapts to sustain cardiac function. While traditional 2D ultrasound measures indicated that the pregnant heart can adapt to improve functional performance in the presence of pressure overload, more advanced 4D strain measures showed signs of early cardiac decline. This underscores the need for a multi-scale assessment of left ventricular health in pregnant patients with hypertension to ensure that cardiac function is accurately evaluated and potential dysfunction is appropriately identified.

## Abbreviations

angII: Angiotensin II
ANGII: Angiotensin II-treated
CO: Cardiac output
dSR: Early diastolic strain rate
E/e: Embryonic day
*E_cc_*: Circumferential strain
*E_ll_*: Longitudinal strain
EDV: End diastolic volume
EF: Ejection Fraction
MAP: Mean arterial pressure
P: Postpartum day
PSV: Peak systolic volume
PREG: Pregnant
sSR: Systolic strain rate
SV: Stroke volume
TPR: Total peripheral resistance

## Acknowledgments

We would like to acknowledge Pierre Sicard, PhD for his guidance and feedback in manuscript preparation.

## Data Availability

Source data for this study are openly available at

https://app.box.com/s/jgdsge9ni09d3cku5h4itok6ntesfdsj.

## Author Contributions

- **E. Ghajar-Rahimi:** Conceived and designed research, performed experiments, analyzed data, interpreted results of experiments, prepared figures, drafted manuscript, edited and revised manuscript, approved final version
- **A. Shen:** Analyzed data, interpreted results of experiments, prepared figures, drafted manuscript, edited and revised manuscript, approved final version
- **A. Meeks:** Conceived and designed research, performed experiments, analyzed data, interpreted results of experiments, edited and revised manuscript, approved final version
- **M. Kaissar:** Conceived and designed research, analyzed data, edited and revised manuscript, approved final version
- **S. Faruqi:** Performed experiments, analyzed data, interpreted results, prepared figures, edited and revised manuscript, approved final version
- **S. Stebbings:** Performed experiments, analyzed data, drafted manuscript, edited and revised manuscript, approved final version
- **S. Grev:** Performed experiments, analyzed data, approved final version
- **J. Castro:** Performed experiments, analyzed data, prepared figures, approved final version
- **A. Cox:** Pathological report, edited and revised manuscript, approved final version
- **K. Yoshida:** Conceived and designed research, edited and revised manuscript, approved final version
- **C.J. Goergen:** Conceived and designed research, edited and revised manuscript, approved final version

## Supplemental

**Table S1:** Tail cuff blood pressures, ultrasound-derived cardiac output, and total peripheral resistance. PREG=pregnant, ANGII=angiotensin II treated, e=embryonic day, p=postpartum. \**p<*0.05 vs baseline, †*p<*0.05 vs pregnant cohort, #*p<*0.05 vs ANGII cohort. Values=Mean*±*SEM.

|  | Day | PREG | ANGII | ANGII+PREG |
| --- | --- | --- | --- | --- |
| **Systolic Blood Pressure (mmHg)** | 0 | 103.98 ± 1.33 | 109.22 ± 3.51 | 104.39 ± 2.67 |
|  | 14 | N/A | **138.45 ± 4.35\*** | **125.60 ± 5.26\*** |
|  | 31/e12 | **104.33 ± 3.56#** | **133.92 ± 6.64\*** | **100.65 ± 3.68#** |
|  | 38/p1 | **107.84 ± 2.80#** | **127.32 ± 9.45\*** | 113.52 ± 5.79 |
| **Diastolic Blood Pressure (mmHg)** | 0 | 81.07 ± 2.58 | 82.03 ± 2.88 | 83.77 ± 3.07 |
|  | 14 | N/A | **105.39 ± 4.38\*** | 98.23 ± 5.39 |
|  | 31/e12 | **80.90 ± 3.05#** | **102.59 ± 5.10\*** | **71.25 ± 4.44#** |
|  | 38/p1 | **79.49 ± 2.14#** | **97.56 ± 8.61\*** | 84.51 ± 2.36 |
| **Mean Arterial Pressure (mmHg)** | 0 | 88.38 ± 1.98 | 90.76 ± 3.08 | 90.27 ± 2.86 |
|  | 14 | N/A | **116.06 ± 4.30\*** | **106.99 ± 5.31\*** |
|  | 31/e12 | **88.40 ± 3.05#** | **112.66 ± 5.55\*** | **80.74 ± 4.09#** |
|  | 38/p1 | **88.61 ± 1.93#** | **107.19 ± 8.85\*** | 93.81 ± 3.35 |
| **Cardiac Output (mL/min)** | 0 | 10.75 ± 1.44 | 12.37 ± 0.92 | 9.38 ± 0.97 |
|  | 14 | N/A | 13.00 ± 0.83 | 10.58 ± 0.93 |
|  | 31/e12 | **16.09 ± 1.13#\*** | 11.68 ± 0.98 | **15.33 ± 1.21#\*** |
|  | 38/p1 | **13.67 ± 1.22#** | 9.80 ± 1.33 | **14.51 ± 1.02#\*** |
| **Total Peripheral Resistance (mmHg\*min/mL)** | 0 | 8.75 ± 1.44 | 7.56 ± 0.57 | 10.01 ± 0.96 |
|  | 14 | N/A | 9.14 ± 0.66 | 10.32 ± 0.73 |
|  | 31/e12 | **5.57 ± 0.40#** | 10.17 ± 1.10 | **5.42 ± 0.56#\*** |
|  | 38/p1 | **6.83 ± 0.76#** | **12.31 ± 2.14\*** | **6.61 ± 0.58#** |

**Table S2:** Left ventricle function measured from 2D parasternal long-axis ultrasound. PREG = pregnant, ANGII = angiotensin II treated, e = embryonic day, p = postpartum. \**p <* 0.05 vs baseline, †*p <* 0.05 vs pregnant cohort, #*p <* 0.05 vs ANGII cohort.Values=Mean*±*SEM.

|  | Day | PREG | ANGII | ANGII+PREG |
| --- | --- | --- | --- | --- |
| **End Diastolic Volume (uL)** | 0 | 31.17 ± 2.93 | 35.59 ± 1.98 | 29.82 ± 3.10 |
|  | 25/e6 | 30.55 ± 0.68 | **44.91 ± 2.97\***† | 43.89 ± 4.82 |
|  | 31/e12 | **44.60 ± 1.54\*** | **46.53 ± 2.33\*** | **57.91 ± 4.99\*** |
|  | 34/e15 | 42.76 ± 2.40 | 45.76 ± 2.39 | **59.17 ± 4.66\*** |
|  | 36/e17 | 42.19 ± 6.06 | 45.34 ± 3.87 | 53.68 ± 5.80 |
|  | 38/p1 | 41.32 ± 3.26 | 38.82 ± 4.58 | **50.87 ± 2.12\*** |
| **Peak Systolic Volume (uL)** | 0 | 10.01 ± 0.74 | 13.40 ± 1.48 | 11.20 ± 1.62 |
|  | 25/e6 | 10.59 ± 0.65 | **23.36 ± 2.12†** | 20.64 ± 2.18† |
|  | 31/e12 | 15.41 ± 0.87 | **24.67 ± 3.11\***† | 29.82 ± 3.31 |
|  | 34/e15 | 16.82 ± 2.29 | 24.22 ± 3.46 | 28.82 ± 3.64 |
|  | 36/e17 | 13.71 ± 2.85 | **25.13 ± 2.65†** | 26.00 ± 4.79 |
|  | 38/p1 | 13.97 ± 1.73 | 21.08 ± 3.96 | 20.46 ± 3.08 |
| **Stroke Volume (uL)** | 0 | 21.16 ± 1.35 | 22.95 ± 1.56 | 16.62 ± 2.21 |
|  | 25/e6 | 19.96 ± 0.70 | 21.55 ± 1.96 | 23.29 ± 2.68 |
|  | 31/e12 | 29.19 ± 1.28 | 21.83 ± 2.01 | **28.09 ± 1.93#** |
|  | 34/e15 | 25.90 ± 1.81 | 21.54 ± 2.32 | 30.35 ± 3.29 |
|  | 36/e17 | 28.47 ± 4.47 | 20.21 ± 2.11 | 27.70 ± 1.85 |
|  | 38/p1 | **27.36 ± 2.02#** | **17.74 ± 2.36\*** | **30.42 ± 1.96#** |
| **Ejection Fraction (%)** | 0 | 67.26 ± 2.47 | 63.29 ± 2.95 | 62.77 ± 2.45 |
|  | 25/e6 | 65.34 ± 1.84 | **47.95 ± 3.34†** | **52.93 ± 0.49†** |
|  | 31/e12 | 65.44 ± 1.89 | **47.64 ± 5.00†** | **48.90 ± 1.65†** |
|  | 34/e15 | 61.02 ± 4.49 | 47.90 ± 5.61 | 51.55 ± 4.88 |
|  | 36/e17 | 68.00 ± 5.01 | **44.85 ± 3.27†\*** | 52.52 ± 4.00 |
|  | 38/p1 | 66.39 ± 1.00 | **48.19 ± 5.84†** | 60.29 ± 4.77 |
| **Cardiac Output (mL/min)** | 0 | 10.75 ± 1.44 | 12.37 ± 0.92 | 9.38 ± 0.97 |
|  | 25/e6 | 10.87 ± 0.31 | 11.53 ± 1.07 | 12.73 ± 1.45 |
|  | 31/e12 | 16.09 ± 1.13 | **11.68 ± 0.98†** | **15.33 ± 1.21\*** |
|  | 34/e15 | 13.57 ± 0.95 | 12.05 ± 1.35 | 16.27 ± 1.92 |
|  | 36/e17 | 15.28 ± 2.50 | 11.61 ± 1.24 | 15.11 ± 1.64 |
|  | 38/p1 | 13.67 ± 1.22 | 9.80 ± 1.33 | **14.51 ± 1.02#** |

**Table S3:**
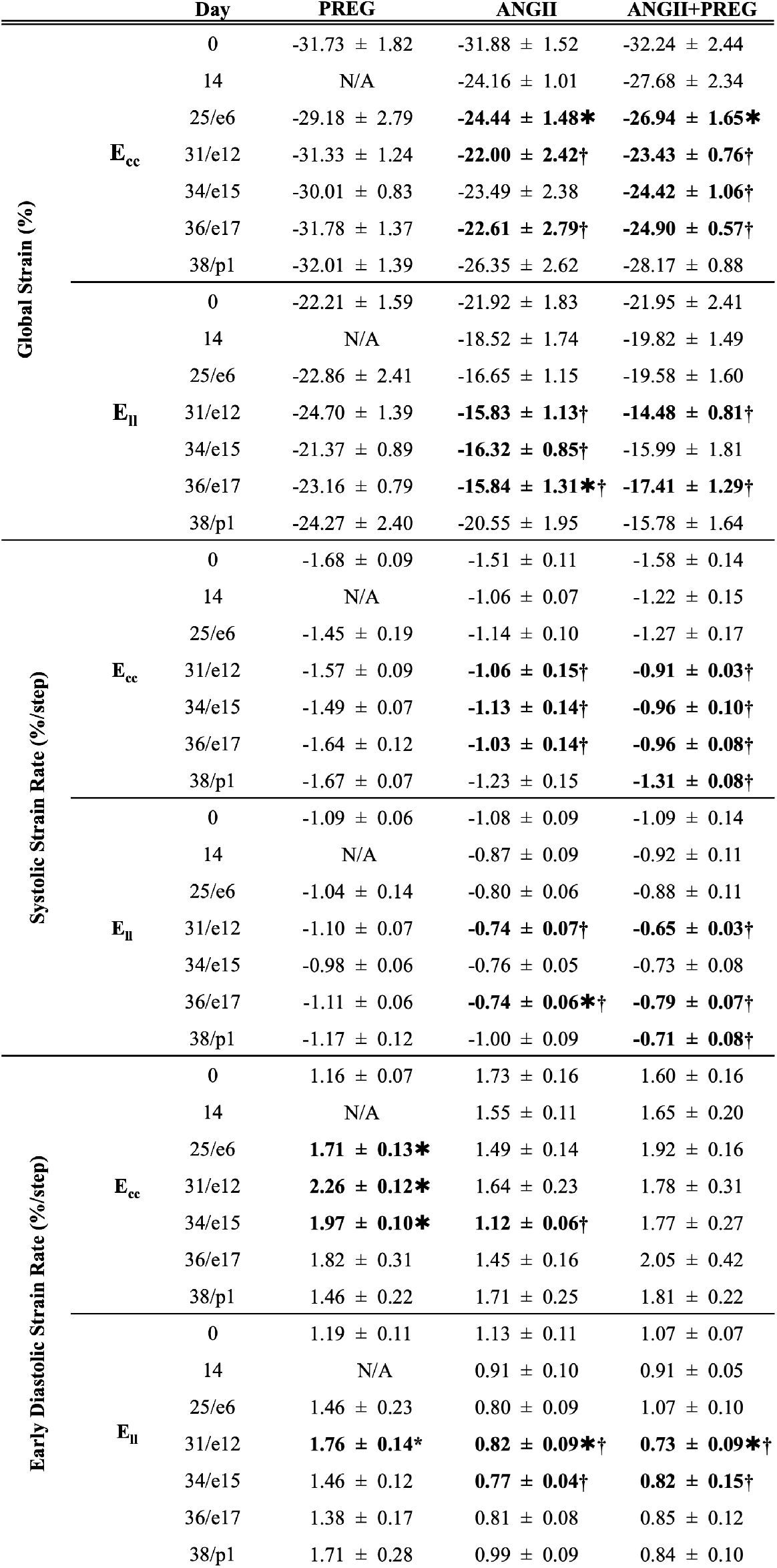
4D ultrasound-derived left ventricular strain, and strain rates for circumferential (*E_cc_*) and longitudinal (*E_ll_*) strain. n=5 for pregnant controls (PREG), n=7 for angiotensin II controls (ANGII), and n=5 for ANGII+PREG. e=embryonic day, p=postpartum. \**p<*0.05 vs baseline, †*p<*0.05 vs pregnant cohort, #*p<*0.05 vs ANGII cohort. Values=Mean*±*SEM.

**Table S4:** Histological Analysis. Lesion presence determined from H&E and MTC histology by a veterinary pathologist. Percent fibrosis measured from MTC in ImageScope, and cross-sectional area measured in ImageJ from WGA. n=5 for pregnant controls (PREG), angiotensin II controls (ANGII) and ANGII+PREG. \**p<*0.05 vs PREG. Values=Mean*±*SEM.

|  | PREG | ANGII | ANGII+PREG |
| --- | --- | --- | --- |
| **Lesion Present (% of animals)** | 0 | 29 | 40 |
| **Percent Fibrosis (% pixels)** | 2.9 ± 0.1 | 5.6 ± 0.6* | 4.8 ± 0.6 |
| **Cross-sectional Area ($\mu\text{m}^2$)** | 199.6 ± 8.9 | 188.0 ± 6.8 | 188.0 ± 8.0 |

**Figure S1:**
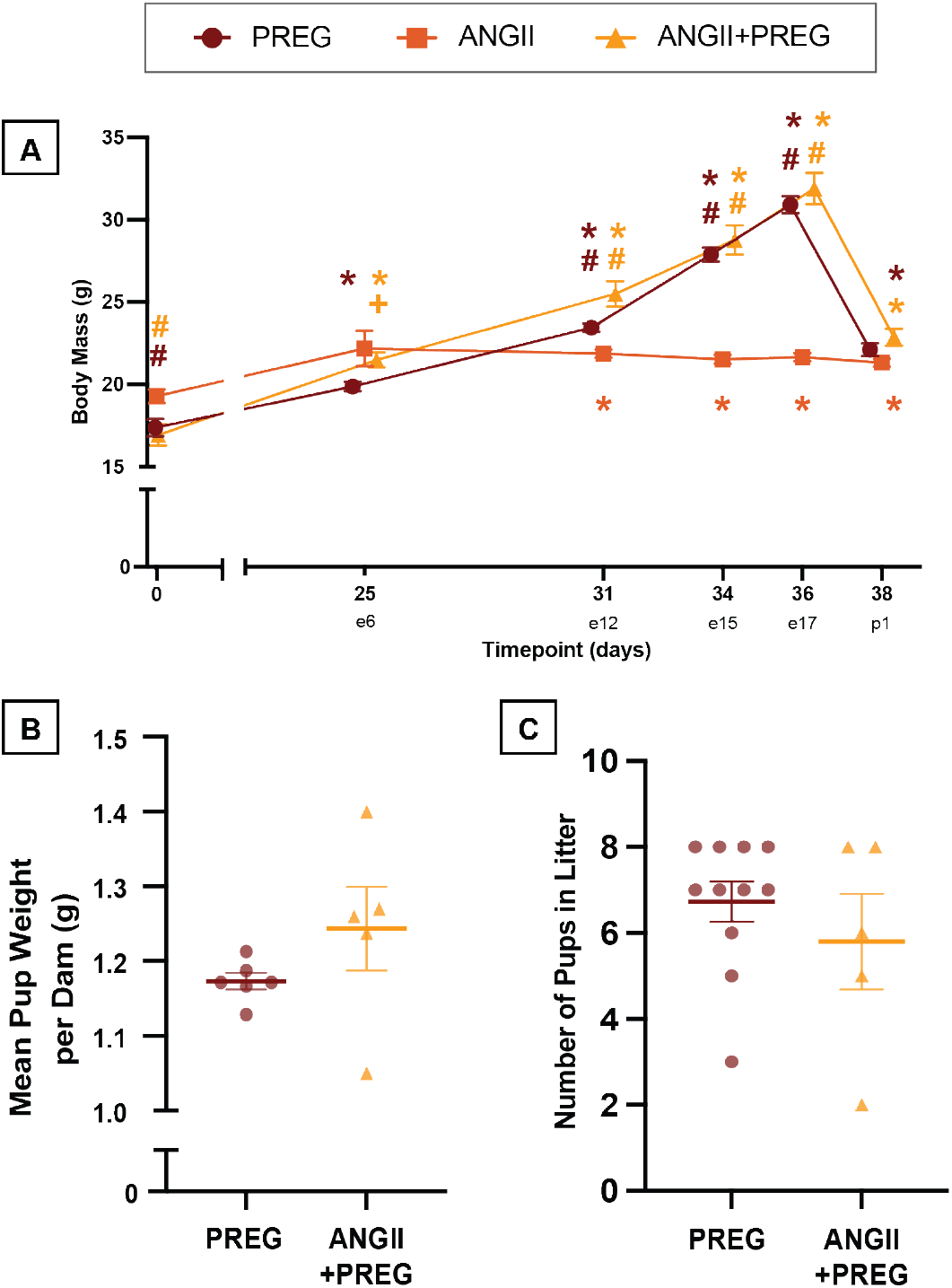
Body weights and litter outcomes. A) Dam weight throughout experimental timeline and B) mean pup weight per dam on postpartum day 1, and C) litter size. n=5 for pregnant controls (PREG), n=7 for angiotensin II controls (ANGII), and n=5 for ANGII+PREG. \**p<*0.05 vs baseline, +*p<*0.05 vs pregnant cohort, #*p<*0.05 vs ANGII cohort. Values=Mean*±*SEM.

**Figure S2:**
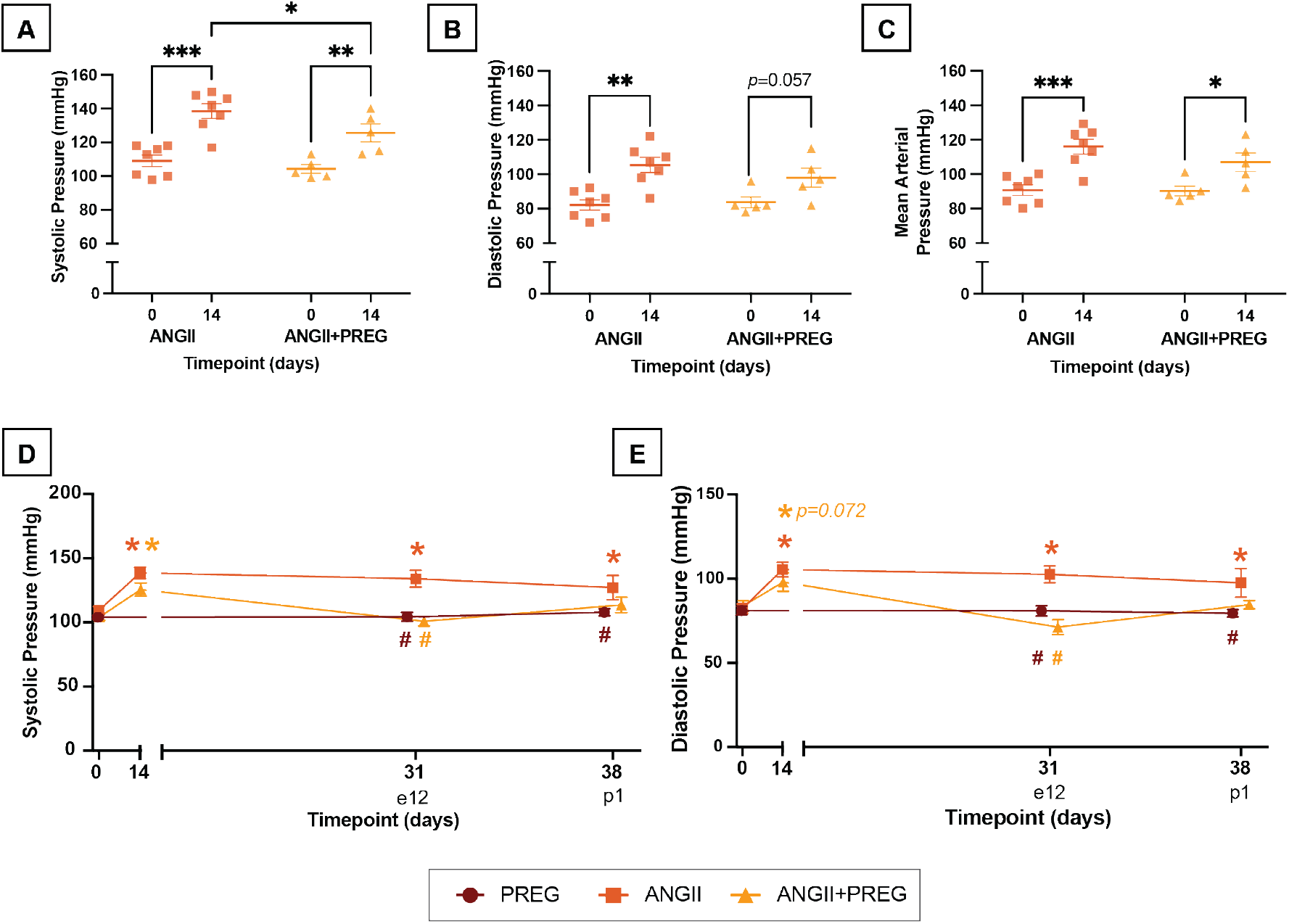
Tail Cuff Blood Pressure. A-C) Systolic, diastolic, and mean arterial pressure (MAP) for angiotensin ii (ANGII) and ANGII+pregnant (ANGII+PREG) groups at baseline and day 14. \**p<* 0.05, \*\**p<* 0.01, \*\*\**p<*0.001 D-E) Systolic and diastolic blood pressure for PREG, ANGII, and ANGII+PREG at baseline, day 31 and 38. \**p<*0.05 vs baseline, +*p<*0.05 vs pregnant cohort, #*p<*0.05 vs ANGII cohort. Color indicates group. n=5 for PREG, n=7 for ANGII, and n=5 for ANGII+PREG. e=embryonic day, p=postpartum day. Values=Mean*±*SEM.

**Figure S3:**
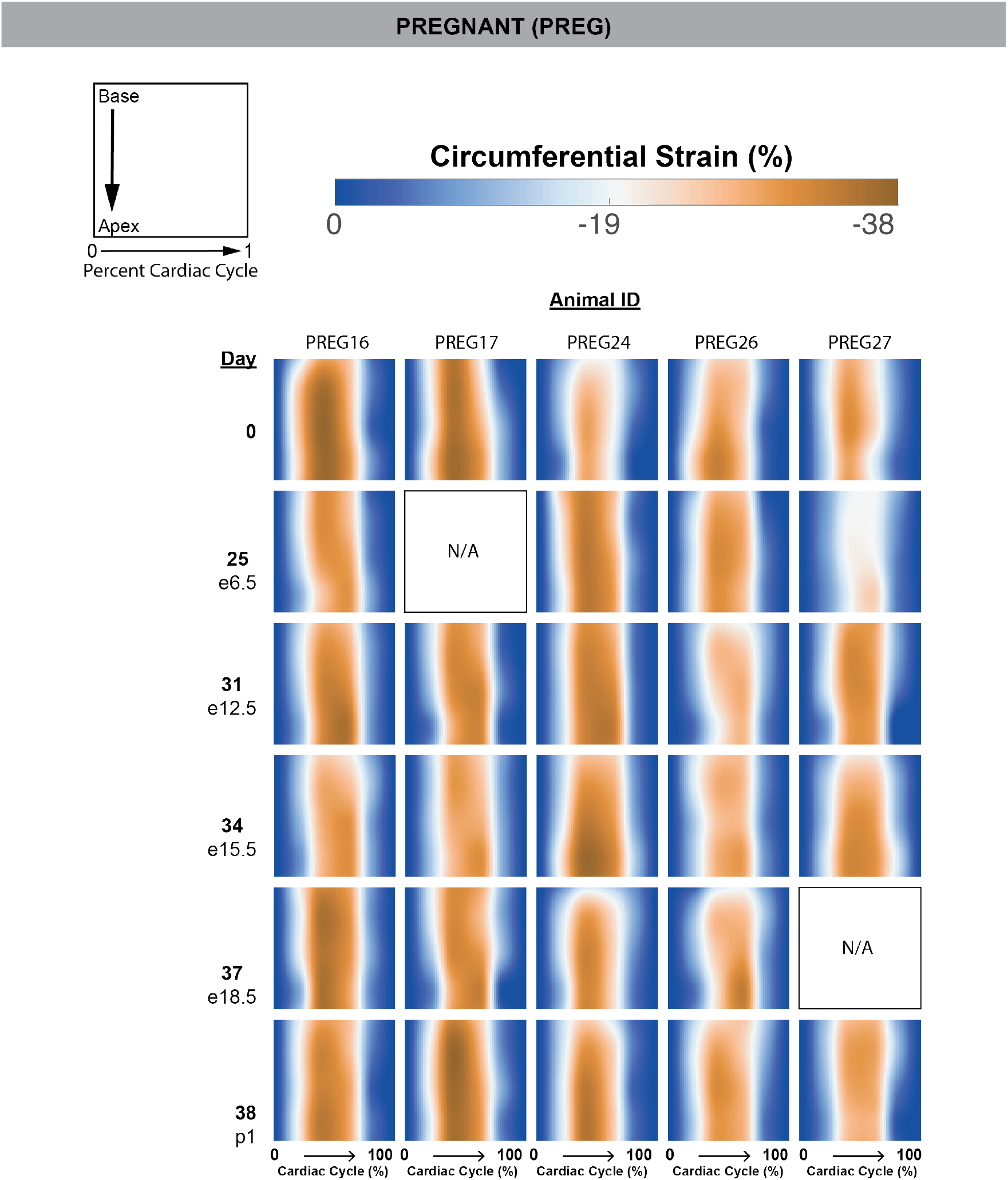
Raw heatmaps of circumferential strain for pregnant animals. e=embryonic day, p=postpartum day.

**Figure S4:**
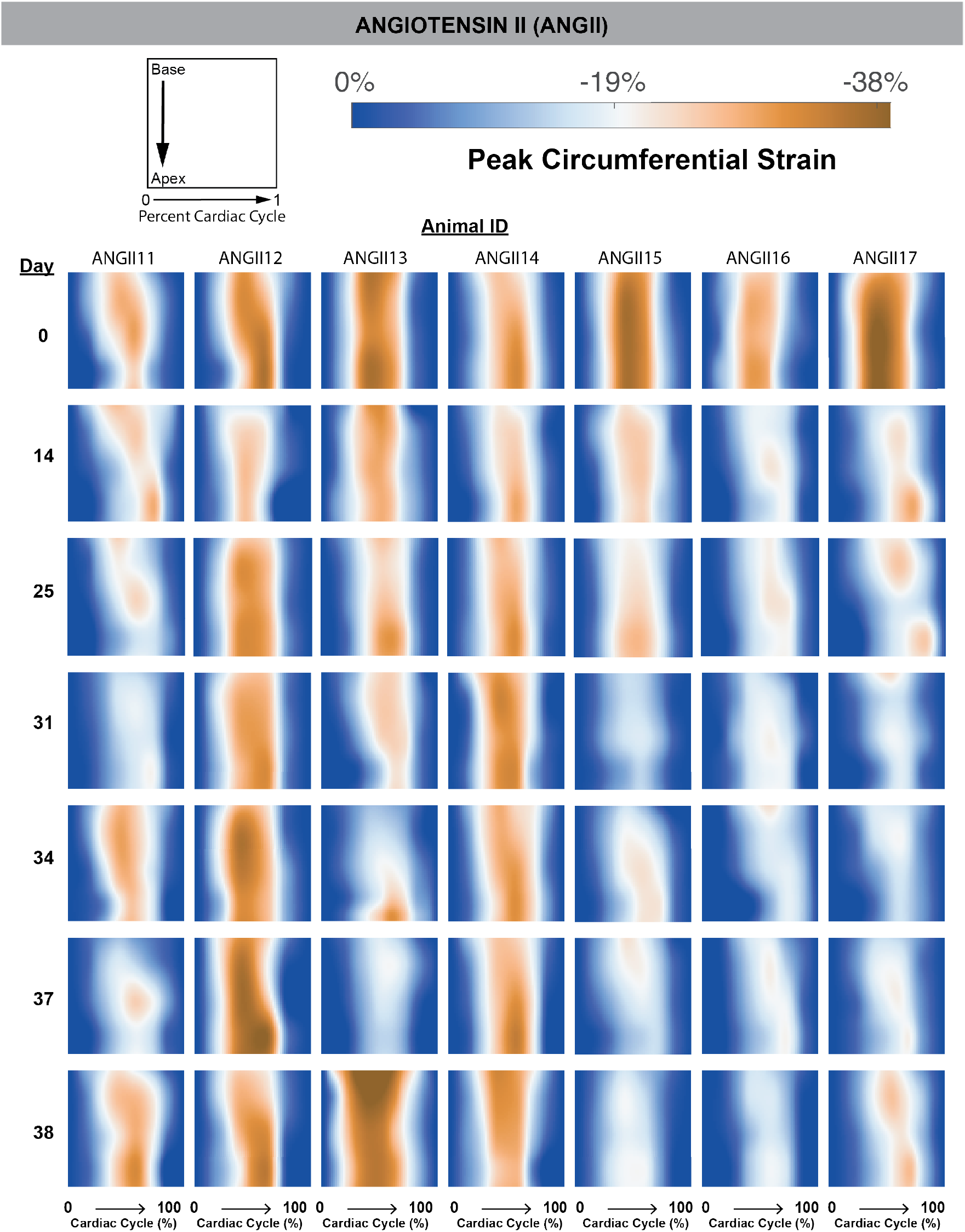
Raw heatmaps of circumferential strain for angiotensin II animals.

**Figure S5:**
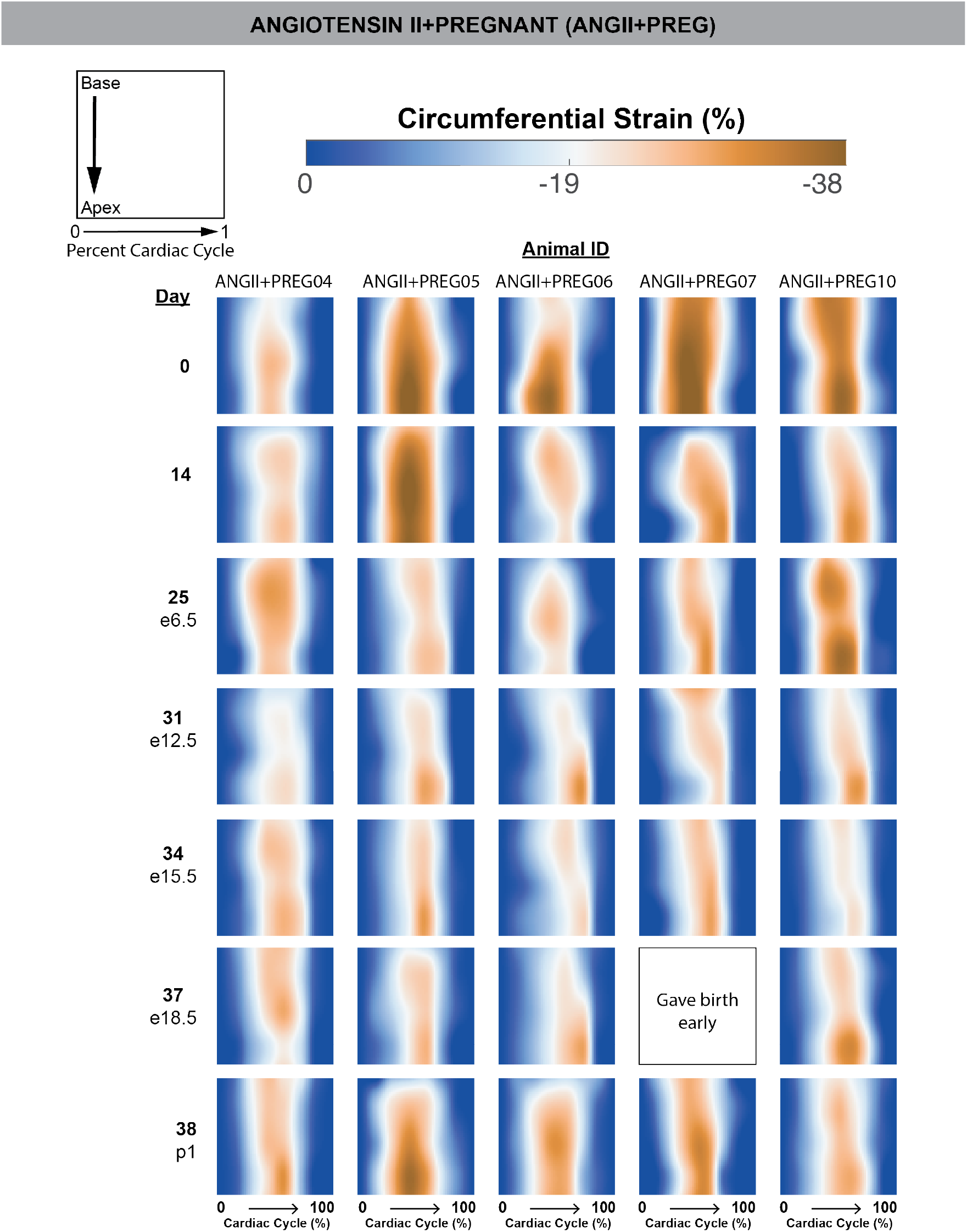
Raw heatmaps of circumferential strain for angiotensin II-pregnant animals. e=embryonic day, p=postpartum day.

**Figure S6:**
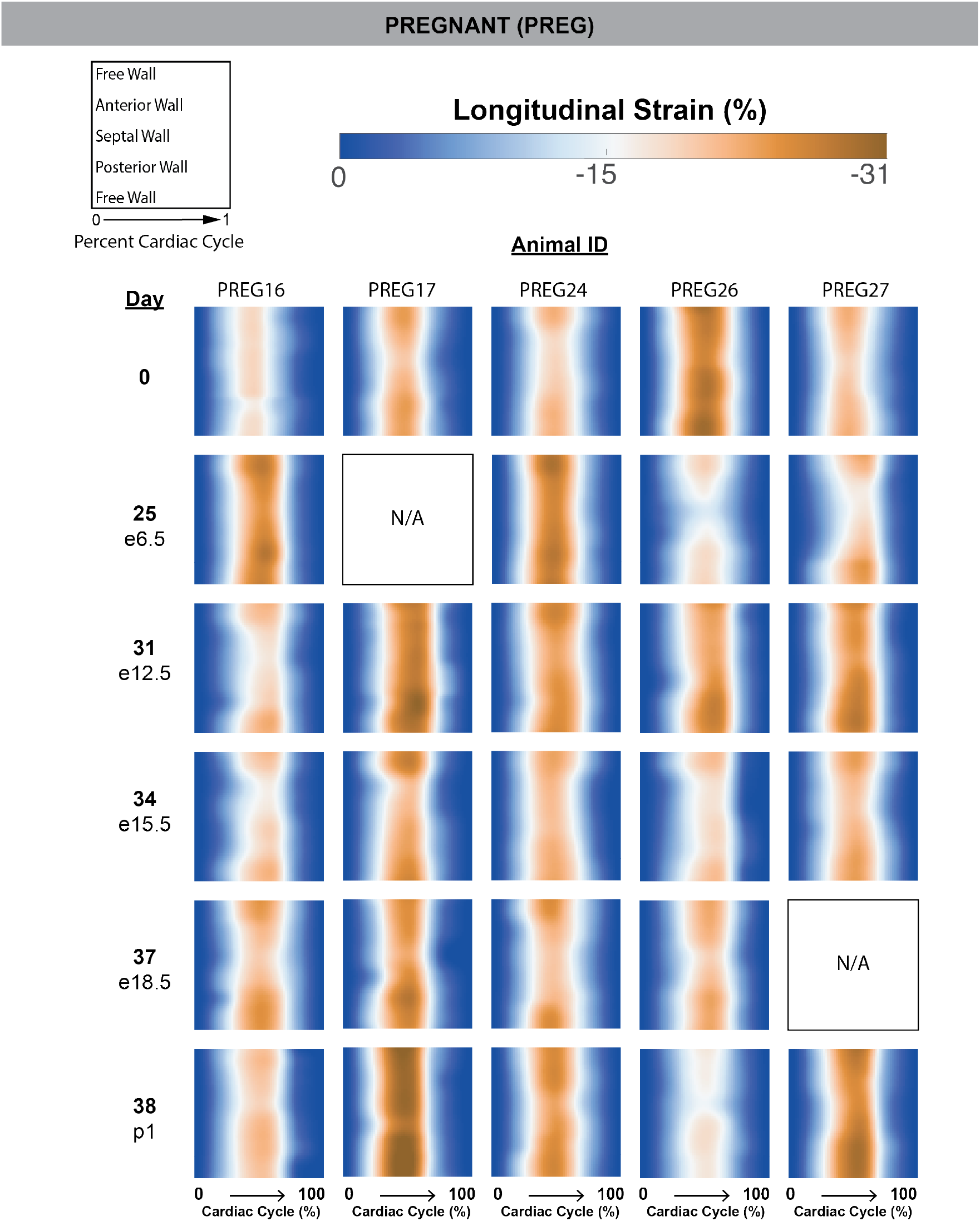
Raw heatmaps of longitudinal strain for pregnant animals. e=embryonic day, p=postpartum day.

**Figure S7:**
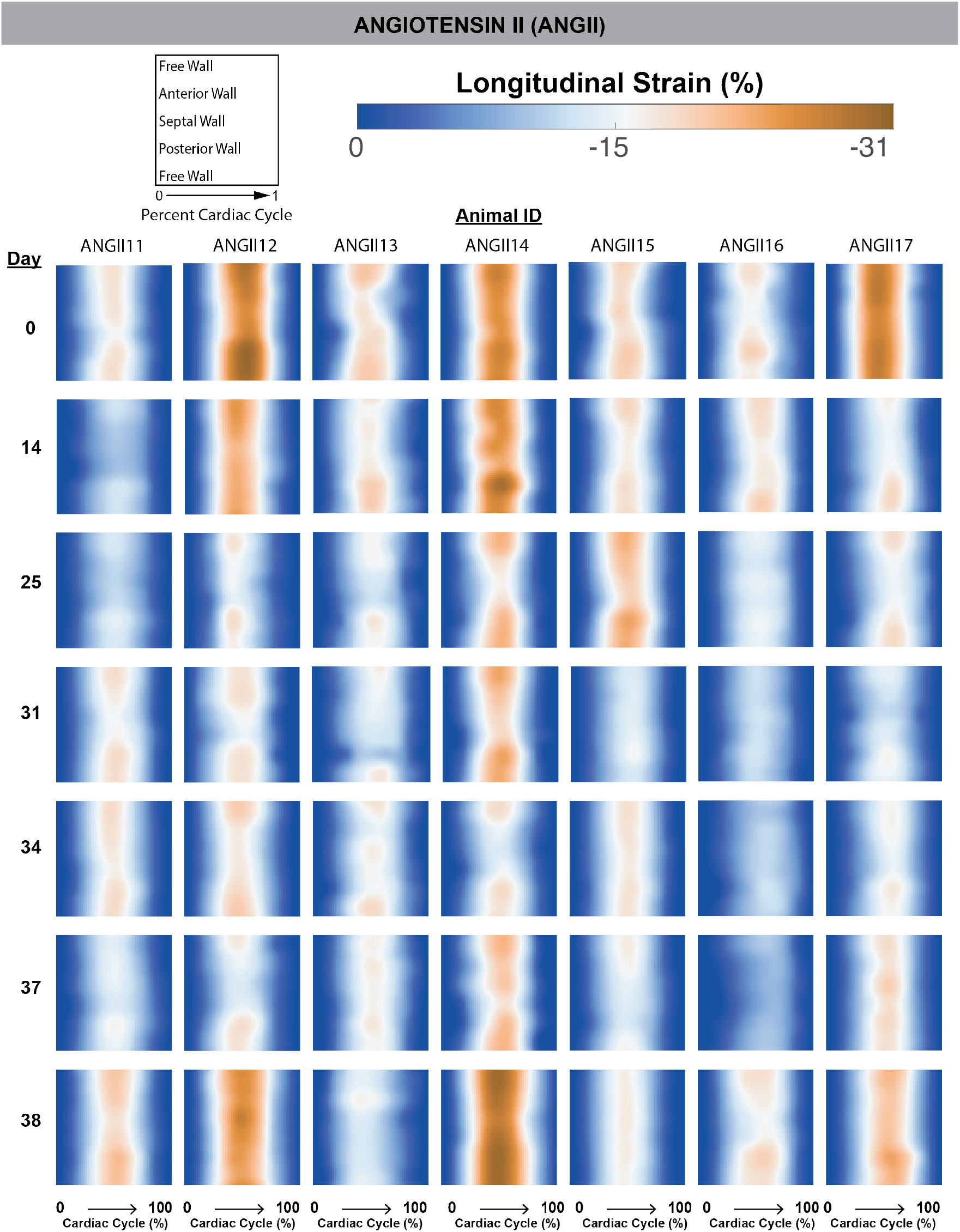
Raw heatmaps of longitudinal strain for angiotensin II animals.

**Figure S8:**
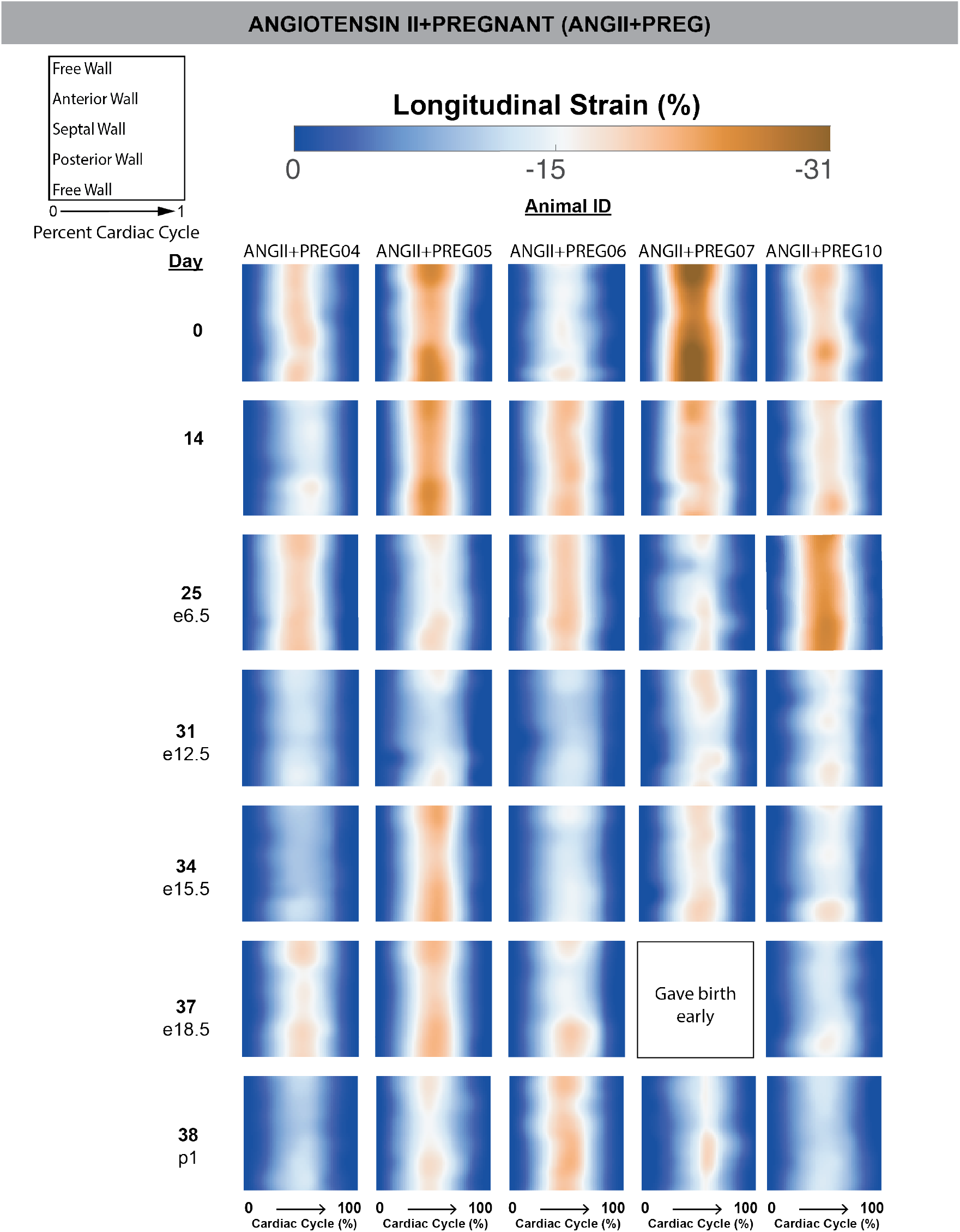
Raw heatmaps of longitudinal strain for angiotensin II treated pregnant animals. e=embryonic day, p=postpartum day.

**Figure S9:**
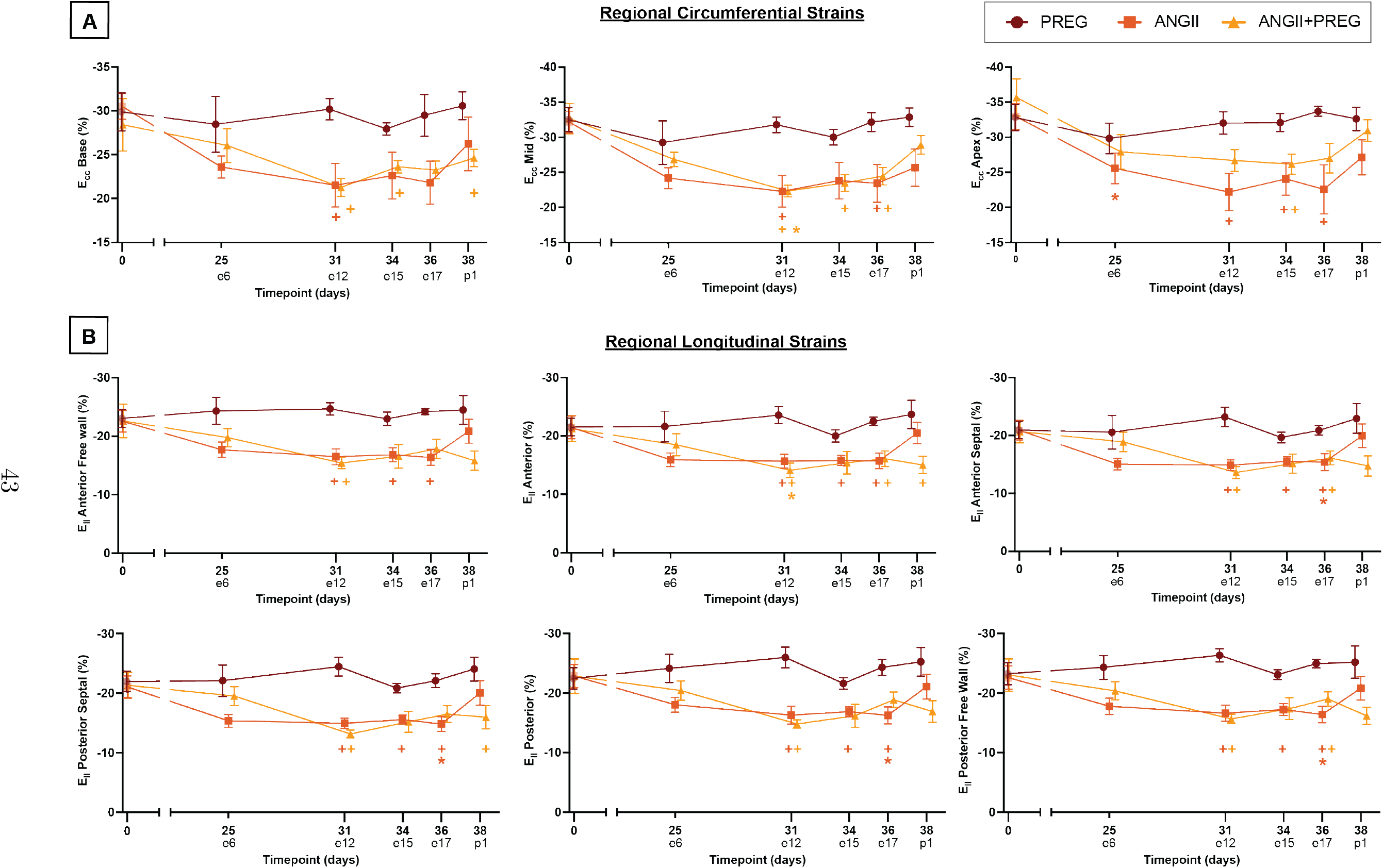
Regional Circumferential and Longitudinal Strains. n=5 for pregnant control (PREG), n=7 for angiotensin II control (ANGII), and n=5 for ANGII+PREG. e=embryonic day, p=postpartum day. Values=mean*±*SEM.

